# Dissociable roles of prefrontal plasticity in decision making strategy and execution of habitual behavior

**DOI:** 10.1101/2025.05.19.654900

**Authors:** Nozomi Asaoka, Diane Pagano, Yasunori Hayashi

**Affiliations:** Department of Pharmacology, Kyoto University Graduate School of Medicine, Kyoto 606-8501, Japan

## Abstract

Habits are essential for sustaining adaptive behaviors but can also lead to maladaptive behaviors such as compulsive and addictive disorders. They emerge through a shift in decision-making from motivation-driven strategy to an automatic one. During this, the level of habit execution, such as frequency and duration, is superficially maintained despite a decline in motivational drive, raising the question of how the amount of execution is maintained even when decision-making strategies undergo substantial changes. By developing a unique paradigm capable of inducing habit formation within a defined time window in male mice, we found that shift in decision-making and amount of habit execution are independently controlled by plasticity in distinct cortical pathways. Erasure of each plasticity selectively altered the decision-making strategy or the execution level without affecting the other. These findings reveal a dual regulatory model for habit, providing new insights into the neurocircuit mechanisms underlying both adaptive and maladaptive habits.

## Introduction

Habits play a crucial role in maintaining beneficial but effortful behaviors, while maladaptive habits cause refractory, over-engaged behaviors shown in compulsive disorders and behavioral addictions^1–5^. Elucidating the regulatory mechanisms of habitual behavior is therefore essential for understanding both adaptive and maladaptive habits^6–8^. Habit formation has been conceptualized as a shift in decision-making—from a flexible, motivation-dependent goal-directed strategy to a more rigid, automatic one^2,5,7^. While this framework captures qualitative differences in behavior selection, it overlooks the quantitative dimension of habitual execution, such as duration and frequency of the behavior, that is critical for evaluating the real-world utility of adaptive habits and the pathological severity of maladaptive ones. Indeed, during the transition, despite the decline in motivational drive, the level of execution is superficially maintained; however, the mechanism that allows the execution level to be preserved despite significant changes in decision-making strategies remains unclear.

This motivated us to propose that the assessment of habitual behavior should incorporate not only the nature of the underlying decision-making strategy but also an additional dimension: regulation of execution level. While decision-making and execution levels in goal-directed behavior are regulated by evaluations of cost and benefit^9,10^, habitual actions, at least at the level of decision-making, proceed automatically without such value-based evaluations. Therefore, a distinct mode of control is likely to govern execution levels in habitual behavior. Based on this premise, we hypothesize the existence of a neural mechanism that stores and maintains the execution level of behavior independently of the mechanisms that govern the transition from goal-directed to habitual strategies.

To address this, we developed a novel behavior task, which allows us to dissect the neuronal mechanisms responsible for the transitions in decision-making strategies together with determining the execution level in the same cohort of animals. Our analyses revealed that the transition in decision-making strategy and execution level are indeed separately controlled. To analyze the circuit mechanism, we combined this task with *ex vivo* slice recordings, *in vivo* Ca^2+^ imaging and a recently-developed optogenetic method to erase long-term potentiation (LTP). We found that predominance of a goal-directed strategy and execution level are orthogonally represented as potentiation of synaptic transmission in different prefrontal cortical regions. By targeting region specific plastic changes, we were able to modify the decision-making strategy without affecting execution level or *vice versa*, supporting the notion that decision strategy and execution level are distinct processes that involve different circuits. Our findings on this dual regulatory model extend the current view of habit regulation and offer new insights for understanding and treating maladaptive habits in disorders such as obsessive-compulsive disorder and addiction.

## Results

### Experimental induction of a transition between goal-directed and habitual behaviors

Previous studies have demonstrated that training with ratio-based and interval-based operant conditioning tasks have contrasting outcomes on animal behaviors^3,5,11^. With ratio-based tasks, where rewards are delivered after a certain number of lever presses, animals acquire goal-directed behavior. In contrast, with interval-based tasks, rewards are delivered on the first lever press after a specific amount of time has passed since the prior reward^12,13^. The number and frequency of lever pressing are less contingent on reward deliveries than in ratio-based tasks and the timing of reward deliveries is less affected by subjects’ lever pressing effort. Owing to these factors, interval-based task designs make it difficult for subjects to understand the association between behavioral effort and reward acquisition. Instead, subjects focus on the behavior that previously led to reward delivery; resulting in the predominance of a habitual over goal-directed strategies^3,14^.

To date, studies have used these two protocols separately to monitor the acquisition of goal-directed and habitual behaviors^3,5^. However, this approach makes identifying the timing of the transition from goal-directed to habitual behaviors difficult, and thus does not allow for the study of the neural mechanisms underlying this transition. Moreover, due to such unclear timing of habit formation, the quantitative changes in action execution accompanied by habit formation could not be adequately analyzed. To overcome these issues, we performed ratio- and interval-based tasks in a sequential fashion in the same cohort of mice and evaluated within-individual transition of decision-making strategy together with that of action execution (Fig. 1A). First, goal-directed behavior was established with a ratio-based task, after which habitual behavior was invoked by switching to an interval-based task, allowing us to track within-subject transitions of decision-making strategy and action execution. Mice were first trained with a continuous reinforcement (CRF) task in an operant chamber for 3 days (days 1–3), where each lever press was rewarded with a drop of sucrose solution from a reward port. Following this, they were trained with a ratio-based, variable ratio (VR) task for 6 days (days 4–9), being rewarded after 10 presses on average on days 4–5, and 20 presses on days 6–9. The lever press rate during VR training increased, indicating successful learning of the action-outcome (lever-reward) association^3,5^ (Fig. 1B). On day 10, the task was seamlessly switched to an interval-based, variable interval (VI) task, where animals were rewarded on the first lever press after a variable time interval from the last reward (ranging from 30 to 90 s, average 60 s). When switched to the VI task, mice slightly but significantly decreased the frequency of lever pressing, compared with animals that continued the VR task (VR+VR group) (Fig. 1B).

**Figure 1.**
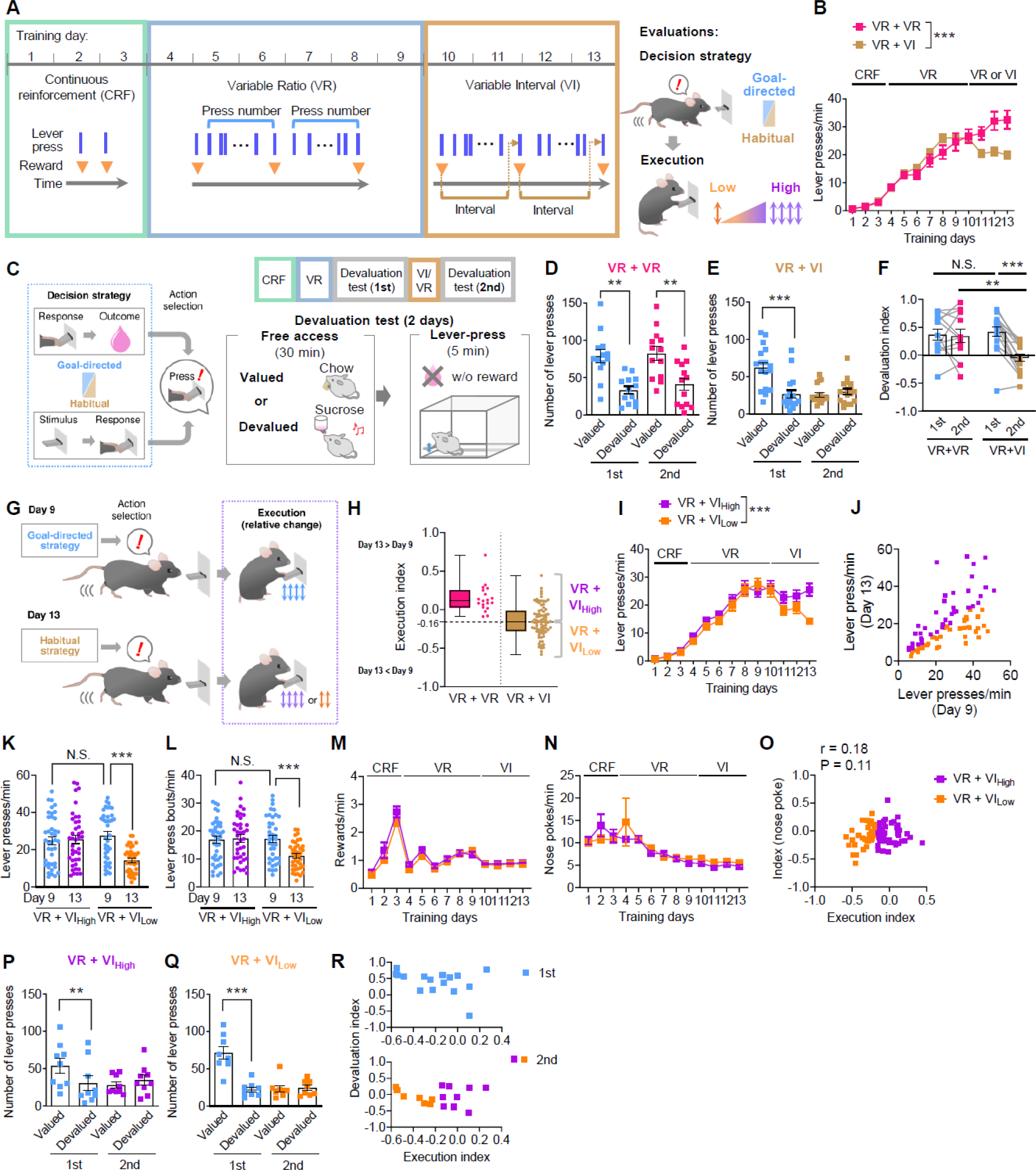
Individual differences in the execution of lever pressing during the two-step training. **(A)** Schematic diagram of the two-step operant training protocol. On day 10, the training schedule was seamlessly switched from a variable ratio (VR) to a variable interval (VI) task (VR+VI). Mice without the VR→VI switching (VR+VR) are considered as the control group. **(B)** Transitions of lever pressing rate in the two-step training. VR+VR, N= 19; VR+VI, N= 79. **(C)** Schematic diagram of the devaluation test for evaluation of decision system. Mice underwent the first devaluation test after the 6^th^ VR session. Then, after the 4^th^ day of VI or VR training, the second devaluation test was performed. On each test day, mice were fed with regular chow, or sucrose solution which mice learned to associate with lever pressing. With free-access to sucrose solution, its relative value declined (devalued condition). After 30-min free-access, mice were placed in the operant chamber and allowed to press the lever for 5 min, during which no sucrose was delivered. **(D, E)** Numbers of lever presses on each test day in the VR+VR (D) and VR+VI (E) groups. **(F)** Within-subject differences in lever pressing between valued and devalued conditions, are shown as the devaluation index. (D–F) VR+VR, N=13; VR+VI, N=17. **(G)** To assess the impact of habit formation on behavioral execution, lever pressing activities before and after habit formation were compared in a within-subject manner. **(H)** Within-subject differences in the lever press rate between days 9 & 13 are plotted as the execution index. The VR+VI group was subdivided into the VR+VI_High_ and VR+VI_Low_ groups by the median value of the execution index. **(I)** Lever press rate in the VR+VI_High_ and VR+VI_Low_ groups. **(J)** Single session lever press rate on days 9 & 13 in the VR+VI_High_ and VR+VI_Low_ groups. **(K)** Comparison of the lever press rate on days 9 & 13, reveal a significant difference in the distribution only on day 13. **(L)** The number of lever press bouts on days 9 & 13. VR→VI switching reduced lever pressing bout in the VR+VI_Low_ group. **(M)** Reward presentation rates in the VR+VI_High_ and VR+VI_Low_ groups. **(N)** Nose poke rates in the VR+VI_High_ and VR+VI_Low_ groups. **(O)** Relations between the execution index and within-subject differences in the nose poke rate between days 9 & 13 are plotted in the XY-plane. (H–O) VR+VI_High_ N=40; VR+VI_Low_ N=39. **(P, Q)** Number of lever presses in each devaluation test in the VR+VI group (E) are separated into the VR+VI_High_ and VR+VI_Low_ groups. (P) N=9, (Q) N=8. **(R)** Within-subject differences in lever pressing between the valued and devalued conditions are shown as the devaluation index. Relations between the execution index and the devaluation index in the first and second devaluation tests are plotted in XY-planes. N=17. **P < 0.01 and ***P < 0.001; N.S., not significant.

To confirm the VR+VI task design led to a transition from a goal-directed to habitual strategy, we conducted two tests: a devaluation test and an omission test^15–17^. In the devaluation test, mice were given free access to either sucrose solution (devalued condition), or regular rodent chow (valued condition) for 30 minutes. They were then subjected to a 5-min lever-press task during which the sucrose reward was not delivered, without signaling this to the mice (Fig. 1C). When mice had already consumed sucrose solution, the sucrose reward is considered “devalued”. In contrast, when mice were pre-fed with rodent chow, the reward remains “valued”. On the following day, the same mice were tested in the opposite condition. When mice employ a goal-directed strategy, they reduce the number of lever presses in the devalued condition compared to the valued condition, as they are already satiated with the sucrose solution. Conversely, when they employ a habitual strategy, the number of lever presses is comparable between the valued and devalued conditions, because the reward value does not affect the decision. Because the absolute number of lever presses during the devaluation test is heavily influenced by the lever press rate during the preceding training, we defined the devaluation index for each mouse, which is the difference in lever pressing between the two conditions, normalized by the total number of lever presses (see Methods). This index gives a positive value when subjects rely on a goal-directed strategy and approaches zero when they employ a habitual strategy, irrespective of absolute lever press number^18^. When the devaluation test was conducted after VR training, there was a significant reduction in lever pressing in the devalued condition resulting in a positive devaluation index (1^st^ test; Fig. 1D–F), confirming that mice employed a goal-directed strategy^5,17^. However, after an additional 4 days of VI training, mice pressed the lever a comparable number of times in both the valued and devalued conditions. This led to devaluation indices close to zero, indicating that mice had changed their strategy to a habitual one with VI training. This was not simply the result of prolonged training, because mice that underwent 13 days of VR training (VR+VR) continued to exhibit a reduction in the number of lever presses in the devalued condition (2^nd^ test; Fig. 1D & F). The effects of devaluation in the VR+VI mice were comparable to those of mice trained for 13 days with the VI paradigm (VI+VI mice. Fig. S1A–C).

In contrast to lever pressing, significant decreases in the number of nose pokes to the reward delivery port were observed in the devalued condition for all groups, except for the first test of the VI+VI group, which exhibited a nonsignificant trend toward reduction. These results suggest that their motivation towards sucrose rewards was reduced irrespective of the training conditions (Fig. S1D–G). In addition, the sucrose/chow intake balance during free access did not differ between conditions (Fig. S1H). Therefore, the lack of inhibitory effects of devaluation on lever pressing in the VR+VI mice was not due to impaired satiety-mediated outcome devaluation, but rather due to their habitual decision strategy, which is not motivated by the current value of sucrose rewards. Consequently, these results confirm that VR→VI switching, but not extended VR+VR training leads to the formation of habitual behavior.

Habit formation was also assessed in the omission test^17^. In the task conducted on the day after the completion of VR+VI training, sucrose solution was delivered automatically every 20 seconds when the lever was not pressed, whereas lever pressing triggered a delay in reward delivery, thereby degrading the association between lever pressing and sucrose delivery (Fig. S1I). Mice utilising a goal-directed strategy diminish the number of lever presses more quickly than those adopting a habitual strategy^17^. Consistent with this idea, the number of lever presses decreased slower in the VR+VI group, compared with the VR+VR group (Fig. S1J & K).

Taken together, these data indicate that implementing a VI training period, following VR training (VR+VI task) converts an established goal-directed behavior into habitual behavior, thereby providing a versatile paradigm to allow for the detailed analysis of the regulatory mechanisms underlying newly formed habitual behaviors.

### Execution level of habitual behavior is controlled independently from the decision-making strategy

Our VR+VI task data revealed there was considerable variation within-subjects in lever pressing changes associated with the transition from a goal-directed to habitual decision strategy (Fig. 1G–O). We hypothesized that this variation may prove valuable for identifying the neuronal mechanisms that determine execution level. To this end, we calculated an execution index for each mouse by normalizing the difference in the frequency of lever presses between days 9 (final VR session) and 13 (final VI session), by the total of the frequencies in the two sessions (Fig. 1G, Methods). The execution index varied among mice trained in the VR+VI task from positive to negative with a median value of -0.16 (max: 0.44, min: -0.58, standard deviation (SD): 0.21). In contrast, mice trained with the VR+VR protocol showed significantly higher execution indices with a median of 0.11 (max: 0.71, min: -0.09, SD: 0.18), indicating that the majority of mice maintained or increased their lever pressing rate after 4 days of additional VR training (Fig. 1H).

Next, for further analysis, the VR+VI group was operationally subdivided into two groups; VR+VI_High_ and VR+VI_Low_, based on execution indices falling above or below the median value (-0.16) (Fig. 1H). The lever press rate on day 9 was comparable between the VR+VI_High_ and VR+VI_Low_ groups and did not affect this classification (Fig. 1I–K). Likewise, the number of lever press bouts (see Methods for definition) was comparable on day 9 between the VR+VI_High_ and VR+VI_Low_ groups, however, on day 13, it was significantly decreased in the VR+VI_Low_ group (Fig. 1L). These data indicate that there was no significant difference in behavior between the VR+VI_High_ and VR+VI_Low_ groups during goal-directed behavior, prior to the VR→VI switching. As for other behavioral metrics, there was no significant difference in the frequencies of rewards acquisitions nor nose pokes between the VR+VI_High_ and VR+VI_Low_ groups (Fig. 1M & N). Although there were slight decreases in nose poke frequencies over the course of the training period, the index of nose poke frequency calculated using the same method as the execution index showed no significant correlation with the execution index (Fig. 1O). These observations indicate that differences in execution index between the VR+VI_High_ and VR+VI_Low_ groups resulted from habit formation, rather than reflecting pre-existing variability during goal-directed behavior nor other variations in reward-consuming behaviors.

To test if the strength of habitual strategy correlates with the variation observed in execution indices, we compared the results of the devaluation and omission tests between the VR+VI_High_ and VR+VI_Low_ groups. In the devaluation test, both groups displayed a significant inhibitory effect before VR→VI switching, but not after VR→VI switching (Fig. 1P & Q). Both before and after switching, there was no significant correlation between the devaluation index and execution index (Fig. 1R). Also in the omission test, the decay in lever pressing was comparable across the VR+VI_High_ and VR+VI_Low_ groups, both of which exhibited a slower decay than the VR+VR group (Fig. S1L–N). These data are consistent with the idea that the strength of the habitual strategy does not correlate with execution level.

We next tested if the variation in execution index arises from other behavioral factors by using a T-maze spatial discrimination and reversal learning test and marble burying test, which can measure learning ability, cognitive flexibility and compulsive-like behavior, respectively, after completion of the VR+VI task (Fig. S1O & P). In all experiments, there was no significant difference between the VR+VI_High_ and VR+VI_Low_ groups, suggesting that the differences in execution indices were determined by the daily training, rather than the individual’s general characteristics of cognition or decision-making.

Collectively, these results suggest that the VR→VI switching protocol induces a transition from a goal-directed strategy to a habitual strategy. However, the bias toward habitual strategy was not correlated with behavioral execution level, indicating that they are likely governed by independent mechanisms.

### Cortical region-specific potentiation of synaptic responses in specific task stages

Since repeated training was required for both behavioral learning and habit formation, we hypothesized that training induces plastic changes in synaptic transmission. To search for evidence that plastic changes were responsible for deciding strategy and execution level, excitatory postsynaptic currents (EPSCs) were recorded *ex vivo* from pyramidal neurons in the prefrontal cortical regions. We focused on anterior cingulate, lateral and medial orbitofrontal cortices (ACC, LOFC and MOFC), as they have been implicated in reinforcement learning and decision-making^10,19^. Mice were sacrificed after CRF (day 3), VR (day 9) and VR+VR or VR+VI (day 13) training, after which AMPA/NMDA ratios were recorded from pyramidal neurons in layer 2/3 (L2/3) and layer 5 (L5), to assess the induction of synaptic potentiation of excitatory synaptic transmission^20,21^ (Fig. 2A–L).

**Figure 2.**
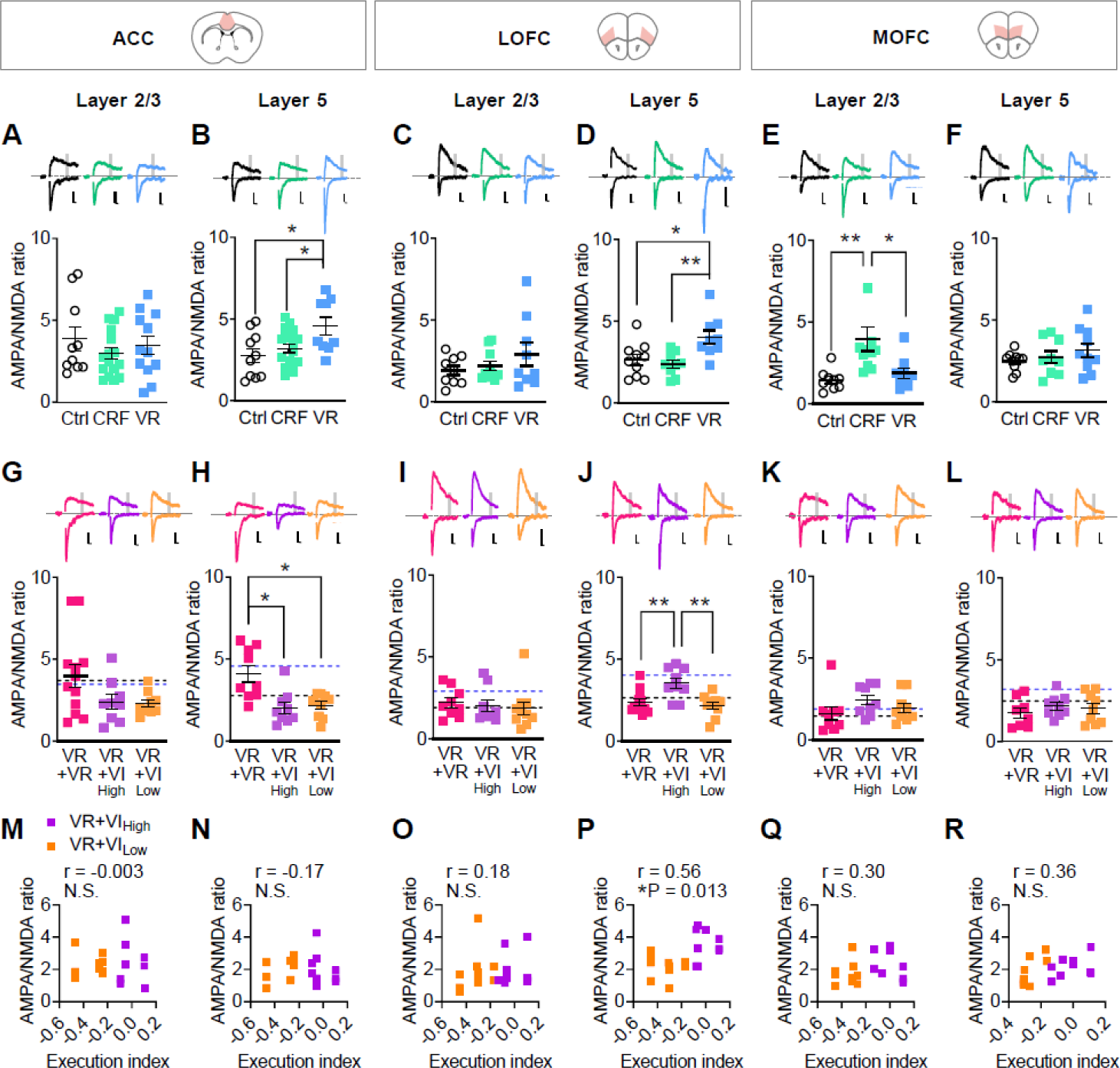
Cortical region-specific plasticity is induced at each training stage. **(A–F)** Acute brain slices of the anterior cingulate cortex (ACC), lateral orbitofrontal cortex (LOFC) and medial OFC (MOFC) were prepared within 30 min after the 3^rd^ CRF and 6^th^ VR training. Control (Ctrl) mice received food restriction but were not trained. AMPA/NMDA ratios were recorded from pyramidal neurons in the ACC layer 2/3 (A), ACC layer 5 (B), LOFC layer 2/3 (C), LOFC layer 5 (D), MOFC layer 2/3 (E) and MOFC layer 5 (F). (A) N=10–16; (B) N=9–18; (C) N=9–10; (D) N=9–10; (E) N=9–11; (F) N=9–10 for each group. **(G–L)** Acute brain slices of the ACC, LOFC and medial OFC were prepared within 30 min after the 4^th^ VI training session. The VR+VR group was introduced as a control matched for total training days. AMPA/NMDA ratios were recorded from pyramidal neurons in the ACC layer 2/3 (G), ACC layer 5 (H), LOFC layer 2/3 (I), LOFC layer 5 (J), MOFC layer 2/3 (K) and MOFC layer 5 (L). The black and blue dashed lines show the mean value of the Ctrl and VR groups in (A–F) respectively. (G) N=9–12; (H) N=9; (I) N=9–10; (J) N=9–12; (K) N=9; (L) N=8–9 for each group. **(M–R)** Correlation between the execution index and AMPA/NMDA ratios in the VR+VI_High_ and VR+VI_Low_ groups. N=18–19. *P < 0.05; **P < 0.01; N.S., not significant.

Following CRF training, there was a specific increase in AMPA/NMDA ratio in MOFC L2/3 neurons, but not in other regions or layers (Fig. 2A–F). However, in the group that underwent subsequent VR training, this increase was diminished. Instead, there were increases in the AMPA/NMDA ratios in both L5 neurons of the ACC and LOFC (Fig. 2B & D). This suggests that as goal-directed behavior is progressively acquired, synaptic potentiation begins in MOFC L2/3 neurons, but transitions to encompass primarily the ACC and LOFC L5 neurons.

Following VR→VI switching (Fig. 2G–L), the AMPA/NMDA ratio in ACC L5 neurons returned to control levels in both the VR+VI_High_ and VR+VI_Low_ groups, whereas it remained elevated in the VR+VR group (Fig. 2G & H). However, the VR+VI_High_ group showed significantly higher AMPA/NMDA values in the LOFC L5 neurons, but not L2/3, compared with both the VR+VR and VR+VI_Low_ groups (Fig. 2I & J). In the MOFC, there were no significant differences in AMPA/NMDA ratios among groups (Fig. 2K & L). Finally, when comparing individual task performance, we observed a significant correlation between the AMPA/NMDA ratio of LOFC L5 neurons and execution index (Fig. 2M–R).

To confirm the findings observed with evoked EPSCs, we recorded Sr^2+^-induced AMPAR-mediated asynchronous EPSCs (aEPSCs) (Fig. S2A–F). The results were consistent with the evoked EPSC data, except for a high aEPSC amplitude observed from LOFC L5 neurons in the VR+VR group (Fig. S2F), which contradict the evoked EPSCs that were comparable to control levels. In the late stages of LTP, NMDA currents are potentiated following potentiation of AMPA currents, resulting in a reset of the AMPA/NMDA ratio^21^. Therefore, it is possible that NMDAR inputs to those LOFC L5 neurons had also been potentiated by prolonged VR+VR training, potentially leading to an overall decrease in AMPA/NMDA ratios, compared with the VR group.

We also tested for changes in presynaptic release probability using the paired-pulse ratio (PPR; Fig. S2G–R). There were no significant differences in the PPR in all regions tested, except for a significant decrease in ACC L5 pyramidal neurons of the VR group, suggesting that presynaptic release probability is also increased.

Together, our observations suggest that region-specific plastic changes in the cortex potentially underlie transitions between goal-directed and habitual strategies, as well as modulating execution levels. Acquisition of a goal-directed behavior potentiated AMPAR-mediated synaptic transmission in the ACC and LOFC L5 neurons. In the ACC L5 neurons, these synaptic inputs were subsequently depotentiated by habit formation. In contrast, execution correlated with potentiation of LOFC L5 neurons, but not other regions tested.

### Chemogenetic inhibition of ACC facilitates habit formation while excitation reverts habitual to goal-directed behavior

To test if the increased excitability of ACC neurons was required for the predominance of goal-directed over a habitual strategy, we employed a chemogenetic approach. The inhibitory receptor hM4Di was expressed in ACC and mice were subsequently trained in the VR task for acquisition of a goal-directed behavior. Next, mice were administered with either vehicle or clozapine-*N*-oxide (CNO) to inhibit ACC neurons prior to the devaluation test (Fig. 3A–D). After the first devaluation test, mice were re-trained with the VR protocol for a day, then underwent a second devaluation test with the reverse drug condition. Lever pressing in mice that received the vehicle was significantly decreased in the devalued condition, however the CNO treated group showed no significant changes between the valued and devalued conditions, indicative of a predominantly habitual strategy. Conversely, when the activity of ACC neurons was increased with hM3Dq after the transition to habitual behavior by the VR+VI task, devaluation induced a significant reduction in lever pressing in the devaluation test, indicating that habitual behavior could be reverted to a goal-directed strategy via artificial ACC modulation (Fig. 3E–H). Additionally, similar effects of ACC activation on the devaluation test were consistently observed in mice trained only with the VI task for 6 days, suggesting that the contribution of ACC in promoting a goal-directed strategy remains irrespective of the training paradigm used (Fig. S3A–C). In all the three experiments, two-way repeated measures ANOVA revealed no significant main effect of CNO administration, suggesting that the effects of chemogenetic manipulations were mainly derived from neural inhibition/activation rather than side effects of CNO (Table S1).

**Figure 3.**
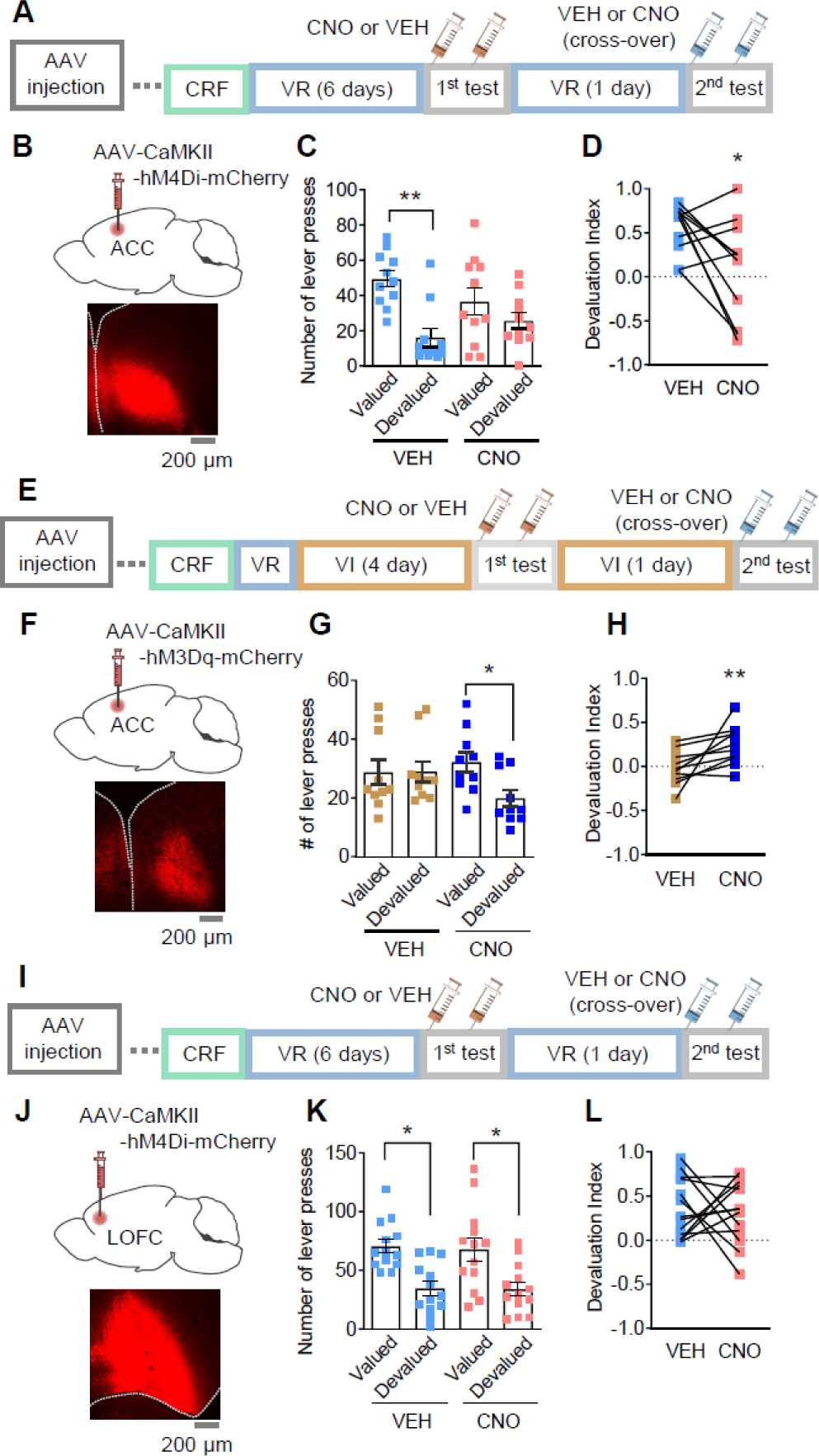
Chemogenetic manipulation of the ACC but not LOFC neurons affects habitual decision-making. **(A)** Mice received injection with AAV-CaMKII-hM4Di-mCherry in the ACC. On the next two days after the 6^th^ VR training, decision-making strategy was assessed by the devaluation test (first test). Before the free-access sessions, clozapine-N-oxide (CNO; 0.5 mg/kg) or its vehicle (VEH) was injected. After the first devaluation test, mice were re-trained with the VR schedule for one day. Then the second devaluation test was conducted with VEH or CNO injection in a cross-over design. **(B)** A representative image of hM4Di-mCherry expression in the ACC. **(C)** Effects of ACC inhibition on lever pressing in the devaluation test. N=11. **(D)** Within-subject changes in the devaluation index of (C). **(E)** Mice received injection with AAV-CaMKII-hM3Dq-mCherry in the ACC. The first devaluation test was performed following the 4^th^ VI training session. After the first test, mice were re-trained with the VI schedule for one additional day, after which the second devaluation test was performed. Same as (A), drug injections were conducted in a cross-over design. **(F)** Representative image of hM3Dq-mCherry expression in the ACC. **(G)** Effects of ACC activation on lever pressing in the devaluation tests. N=10. **(H)** Within-subject changes in the devaluation index of (G). **(I)** Mice received injection with AAV-CaMKII-hM4Di-mCherry in the LOFC. The experimental protocol was same as (A), except for the AAV injection sites. **(J)** Representative image of hM4Di-mCherry expression in the LOFC. N=13. **(K)** Effects of the LOFC inhibition on lever pressing in the devaluation tests. **(L)** Within-subject changes in the devaluation index of (K). *P < 0.05; **P < 0.01.

Interestingly, although VR training induced synaptic potentiation in the LOFC (Fig. 2J & P), chemogenetic inhibition did not affect the devaluation-induced response, unlike ACC inhibition (Fig. 3I–L). Moreover, chemogenetic manipulation of LOFC during the omission test didn’t significantly alter lever pressing when comparing the vehicle- and CNO-treated hM4Di or hM3Dq groups (Fig. S3D–I).

During the chemogenetic experiments, CNO administration did not affect preference or satiety for sucrose rewards in the devaluation tests, as no significant changes were detected in sucrose/chow intake balance, or reduction in nose poking at the reward delivery port in the devalued condition (Fig. S3J–M).

Together, these observations indicate that the suppression of ACC activity can convert behavior from a goal-directed to habitual strategy, whereas activation drives goal-directed behavior. By contrast, the excitability of LOFC neurons does not lead to shifts in decision strategy.

### Projection-specific plasticity of ACC neurons in specific training stages

ACC neurons are a heterogenous population in terms of both gene expression and projection patterns^22,23^. To determine whether training-induced synaptic potentiation occurs in a specific subclass of ACC neurons, which target a particular region, or if it is more global encompassing the entire ACC, we injected retrograde AAV encoding tdTomato into the major targets of ACC: the central striatum (CS), retrosplenial cortex (RSC), and the basolateral amygdala (BLA) to label each subclass of ACC projection neuron^24–26^ (Fig. 4A). Following this, training-induced synaptic potentiation was assessed in tdTomato-positive L5 pyramidal neurons by recording the AMPA/NMDA ratio (Fig. 4B–J). A significant increase in the AMPA/NMDA ratio after VR training was detected only in the RSC-projecting L5 neurons (Fig. 4C). After VR→VI switching, the AMPA/NMDA ratio in this class of neuron returned to control levels, in both the VR+VI_High_ and VR+VI_Low_ groups, whereas the VR+VR group remained elevated (Fig. 4F). We observed no significant correlations between the AMPA/NMDA ratio and execution indices for any subclass of projection neuron tested (Fig. 4H–J). These results indicate that synaptic potentiation in ACC-RSC projection neurons plays an essential role in the acquisition of goal-directed behavior and that its de-potentiation is required for habit formation.

**Figure 4.**
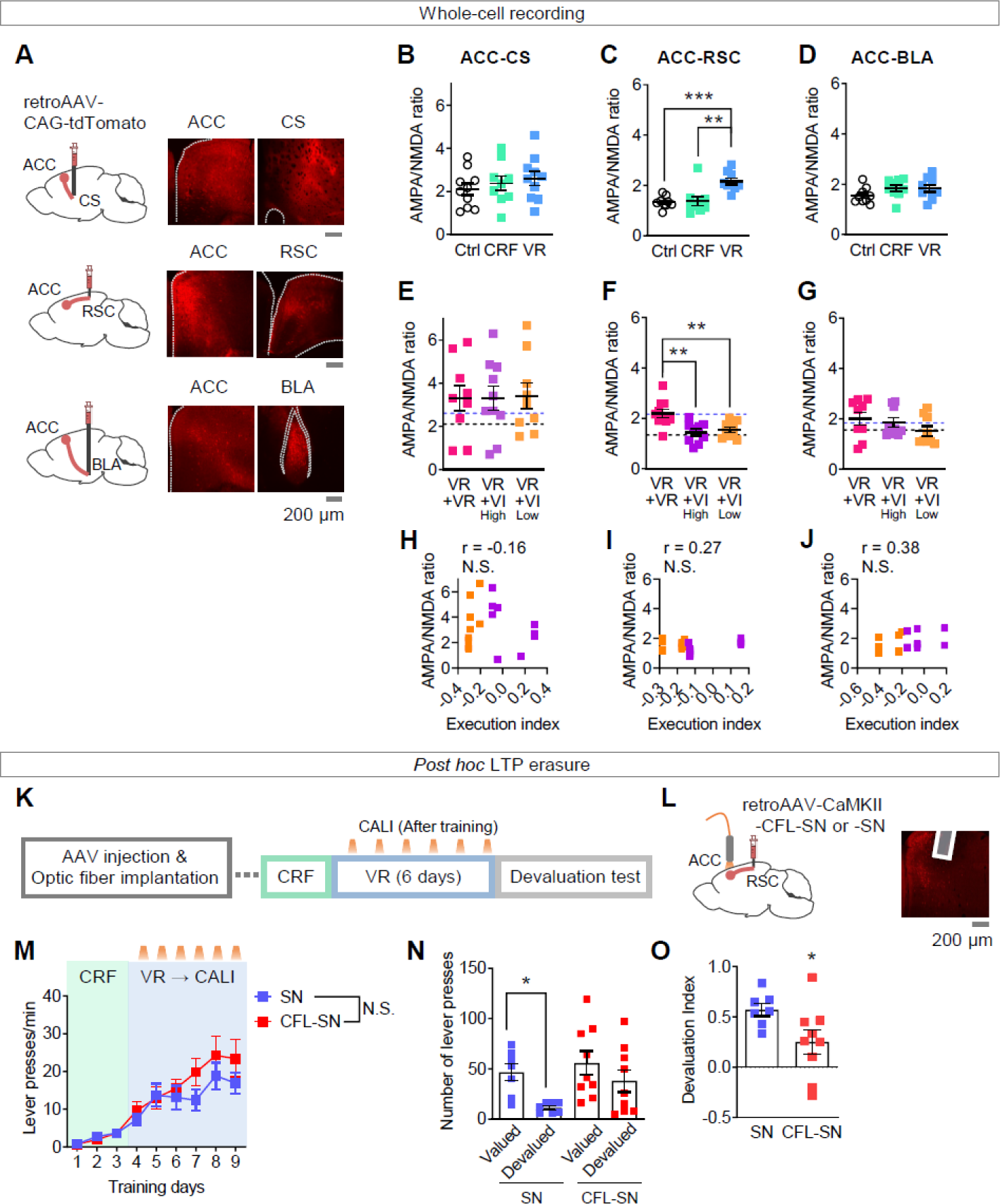
LTP in the RSC-projecting ACC neurons during VR training is necessary for maintenance of goal-directed decision-making. **(A)** Representative images of retrograde labeling with retroAAV-CAG-tdTomato. The retro AAV was injected in the central striatum (CS), retrosplenial cortex (RSC) and basolateral amygdala (BLA). For all three groups, tdTomato-positive neurons were detected in the ACC. **(B–D)** Acute brain slices of ACC were prepared within 30 min after the 3^rd^ CRF and 6^th^ VR training. Control (Ctrl) mice received food restriction but were not trained. AMPA/NMDA ratios were recorded from CS-projecting ACC layer 5 neurons (B), RSC-projecting ACC layer 5 neurons (C) and BLA-projecting ACC layer 5 neurons (D). (B) N=9–10; (C) N=9; (D) N=9 for each group. **(E–G)** Acute brain slices of the ACC were prepared within 30 min after the 4^th^ VI training. The VR+VR group was introduced as a control matched for total training days. AMPA/NMDA ratios were recorded from CS-projecting ACC layer 5 neurons (E), RSC-projecting ACC layer 5 neurons (F) and BLA-projecting ACC layer 5 neurons (G). Black and blue dashed lines show the mean value of Ctrl and VR groups in (B–D) respectively. (E) N=9–10; (F) N=9–10; (G) N=9 for each group. **(H–J)** Correlation between execution index and AMPA/NMDA ratios in the VR+VI_High_ and VR+VI_Low_ groups. N=18–19. **(K)** Schematic diagram of VR training with CALI. Mice received CALI (illumination with 1–2 mW amber light for 1 min) within 5-min after each VR training session ended (days 4–9). **(L)** Representative image of CFL-SN expression and optic fiber implantation in the ACC. retroAAV-CaMKII-CFL-SN or retroAAV-CaMKII-SN were injected into the RSC and optic fibers implanted into the ACC. **(M)** Lever press rate during training with CALI. **(N)** Effects of CALI in RSC-projecting ACC neurons on lever pressing in the devaluation tests. **(O)** Within-subject changes in the devaluation index of (N). SN, N=7; CFL-SN, N=9. *P < 0.05; **P < 0.01; ***P < 0.001; N.S., not significant.

### Optical erasure of synaptic potentiation in RSC-projecting ACC neurons facilitates the transition to habit

To further test this idea, we employed the recently-developed optogenetic tool, cofilin-SuperNova (CFL-SN) to erase long-term potentiation (LTP). Cofilin is an actin-binding protein that accumulates in dendritic spines and stabilizes filamentous actin after the induction of LTP^27^. Inactivation of cofilin by chromophore-assisted light inactivation (CALI) using a photosensitizer protein SuperNova (SN) erases LTP *post hoc,* without affecting basal synaptic transmission or the stability of pre-existing synapses^28^. CFL-SN was expressed in RSC-projecting ACC neurons to determine if *post hoc* LTP erasure would shift the decision strategy from a goal-directed to habitual strategy. To this end, retrograde AAV was used to express CFL-SN, or SN as a control in the RSC-projecting ACC neurons bilaterally (Fig. 4K & L). After each VR session (days 4–9), light was delivered to the ACC through optical fibers to erase training-induced LTP. After a 6-day VR training protocol, a devaluation test was conducted to assess decision strategy. During the VR training, both the CFL-SN and SN groups increased the frequency of lever presses (Fig. 4M), however, in the devaluation test the CFL-SN group showed a comparable number of lever presses in the valued and devalued conditions and a significantly smaller devaluation index than the SN group, indicating that their behavior became habitual (Fig. 4N & O). Since, the two groups had a comparable level of lever pressing during training, this indicates that the acquisition of the lever pressing behavior itself does not require LTP in RSC-projecting ACC neurons (Fig. 4M). Additionally, CALI did not impact the preference and satiety for sucrose rewards in the devaluation processes, suggesting that the evaluation of reward value remains intact (Fig. S4).

Overall, the results suggest that training-induced excitatory synaptic potentiation in the RSC-projecting ACC neurons is required for the maintenance of a goal-directed strategy, but if lost, a habitual strategy becomes predominant. This is consistent with the observation that using VI training to establish habitual behavior, reduced the AMPA/NMDA ratio in this class of neuron (Fig. 4B–J).

### RSC-projecting ACC neurons are active during reward acquisition of goal-directed behavior

Our data highlight the importance of the potentiation of synaptic transmission in a specific class of ACC neuron projecting to RSC, in determining transitions in decision-making strategy. Given their importance and to better understand how these neurons influence behavior, we carried out Ca^2+^-imaging in freely moving animals using a head-mounted miniature fluorescent microscope^29^.

RSC-projecting ACC neurons were targeted by injecting a retrograde AAV vector encoding jGCaMP7s in the RSC (Fig. 5A). A subset of these neurons exhibited burst-like activity during rewarded nose pokes, whereas neurons showed minimal activity during lever pressing or non-rewarded nose pokes (Fig. 5B–D). The mean activity of neurons during rewarded nose pokes was significantly larger during VR training, but reduced following VR→VI switching (Fig. 5E). We classified RSC-projecting ACC neurons into reward-excited and -inhibited cells (Fig. 5F–H; Methods). Significant changes in the fraction of each subclass were observed during the transition from goal-directed to habitual behavior (Fig. 5H). The proportion of reward-excited cells (Fig. 5H) as well as their activity (Fig. 5I) were increased by VR training and decreased following VR→VI switching (Fig. 5I), consistent with our *ex vivo* slice electrophysiology data. In contrast, the proportion of reward-inhibited neurons (Fig. 5H) as well as their reward-evoked inhibition (Fig. 5J) decreased during VR training, but reverted following VR→VI switching. The increase in reward-related excitation followed by goal-directed lever pressing and its reversion during habitual behavior, align with the view that a goal-directed strategy relies on current reward value in decision-making, whereas a habitual strategy makes decisions independent of the value of a reward. There was no significant correlation between changes in activity of these neurons and execution index (Fig. S5). These results confirm that RSC-projecting ACC neurons are selectively potentiated during goal-directed behavior, which reverts upon a transition to habitual behavior.

**Figure 5.**
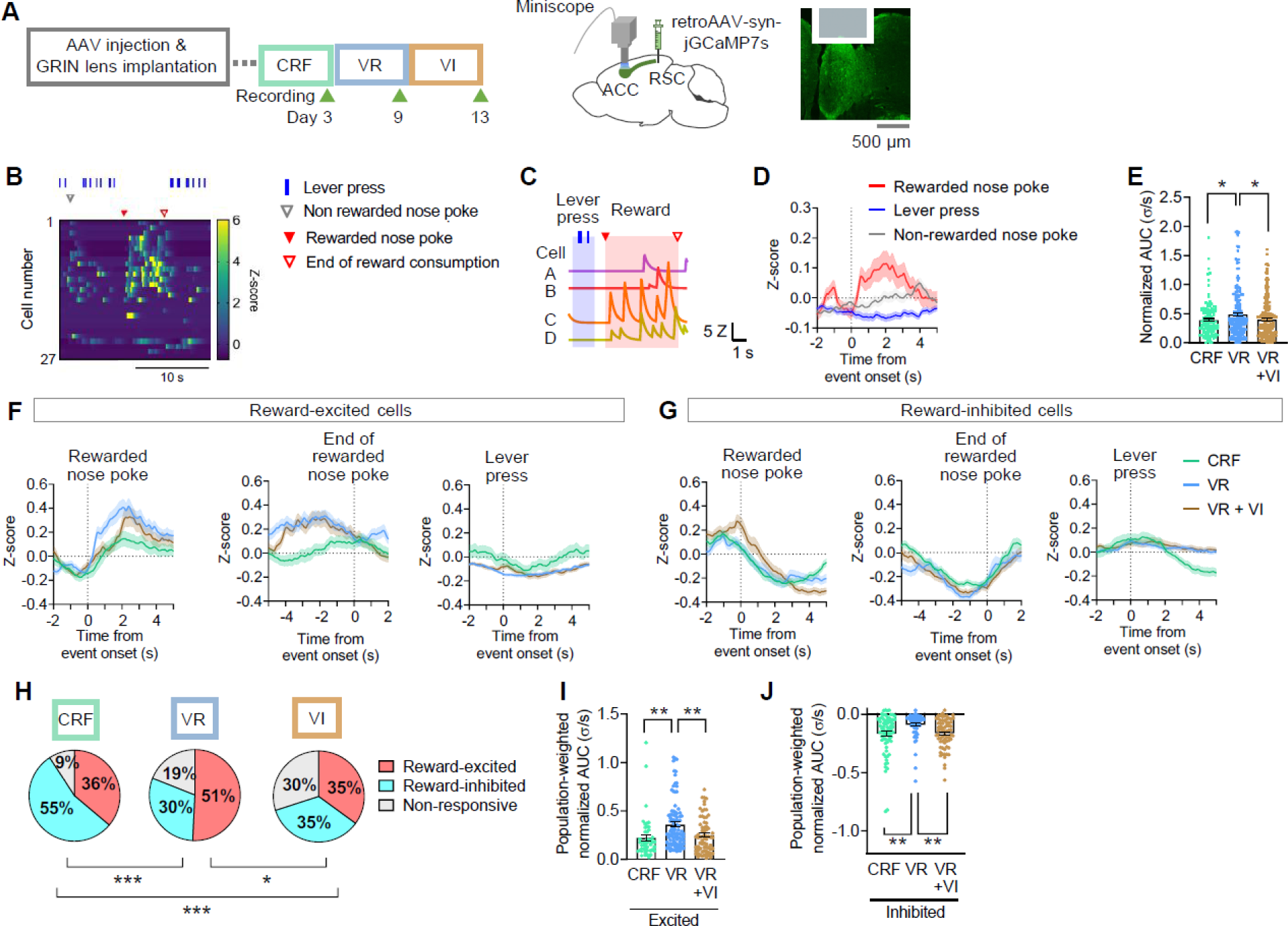
Reward-related activation of RSC-projecting ACC neurons is potentiated during goal-directed action. **(A)** Schematic diagram of *in vivo* Ca^2+^ recording and a representative image of jGCaMP7s expression and GRIN lens implantation. retroAAV-syn-jGCaMP7s was injected into the RSC and GRIN lens was implanted into the ACC. Recordings were performed in the last CRF (day 3), VR (day 9) and VI (day 13) training sessions. **(B)** Representative heat map of Ca^2+^ transients around the first, second (left) and tenth (right) rewarded nose pokes during VR training (day 9). A subset of RSC-projecting ACC neurons were activated during reward consumption. **(C)** The red and blue highlighted zones show the time windows for calculation of reward and lever press-related neural response, respectively (see Methods). **(D)** The average activity of all recorded cells during lever pressing, rewarded nose pokes and non-rewarded nose pokes during VR training. **(E)** The AUC of reward-related responses normalized by the duration of events (from the start to the end of rewarded nose pokes). Each dot represents each cell’s average of normalized AUCs for reward events during training. CRF, N=130; VR, N=195; VR+VI, N=197. **(F,G)** The average temporal dynamics of reward-excited **(F)** and - inhibited **(G)** cells at the start and end of rewarded nose pokes and lever pressing during each training stage. **(H)** Fractions of reward-excited and reward-inhibited cells at each training stage. **(I,J)** For reward-excited and reward-inhibited cells, the population-weighted normalized AUC were calculated by multiplying normalized AUCs of positive (I) or negative (J) reward-related responses by the fraction of the reward-excited and reward-inhibited cells. (I) CRF, N=47; VR, N=99; VR+VI, N=69; (J) CRF, N=71; VR, N=59; VR+VI, N=69. *P < 0.05; **P < 0.01; ***P < 0.001.

### Projection-specific plasticity of LOFC neurons after habit formation correlates with execution index

LOFC layer 5 neurons exhibited an increased AMPA/NMDA ratio during VR training, which was maintained in VR+VI_High_ but not VR+VI_Low_ animals, thereby specifically correlating with execution index after VR→VI switching (Fig. 2J, P). We therefore further explored how LOFC neurons may regulate changes in action execution accompanied by habit formation.

To achieve this, the same retrograde labeling strategy was used to identify specific subclasses of LOFC neurons projecting to CS, RSC, and BLA. We found LOFC projections to both CS and BLA, but not to the RSC (Fig. 6A). The training induced an increase in the AMPA/NMDA ratio, only in the CS, but not BLA-projecting neurons during the VR task (Fig. 6B & C). This ratio remained high in the VR+VI_High_ group, but not in the VR+VI_Low_ group (Fig. 6D & E). Among these two projection populations, only the CS-projecting neurons showed a significant correlation between AMPA/NMDA ratio and the execution index after VR+VI training, suggesting that excitatory transmission of CS-projecting LOFC neurons determines the execution of habitual behavior (Fig. 6F & G).

**Figure 6.**
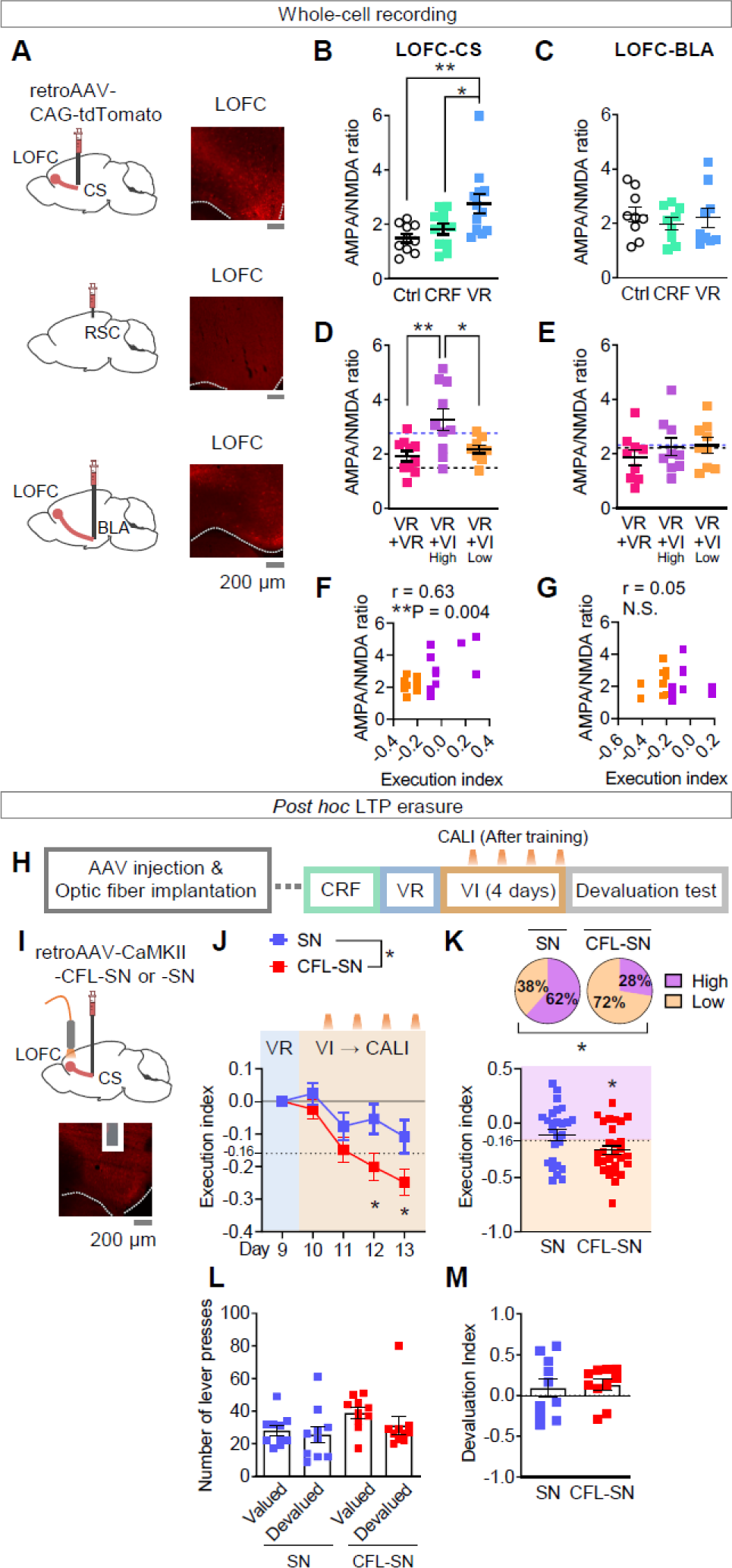
LTP in the CS-projecting LOFC neurons during VI training determined the execution of habitual responses. **(A)** Representative images for retrograde labeling of LOFC neurons with retroAAV-CAG-tdTomato. The retro AAV was injected in the central striatum (CS), retrosplenial cortex (RSC) and basolateral amygdala (BLA). Each image was taken from the same mouse shown in Fig. 4A. tdTomato-positive neurons were not detected in the LOFC, when the retroAAV vector was injected in the RSC. **(B,C)** Acute brain slices of LOFC were prepared within 30 min after the 3^rd^ CRF and 6^th^ VR training. Control (Ctrl) mice underwent food restriction but were not trained. AMPA/NMDA ratios were recorded from CS-projecting LOFC layer 5 neurons (B) and BLA-projecting LOFC layer 5 neurons (C). (B) N=10–12; (C) N=9 for each group. **(D,E)** Acute brain slices of the LOFC were prepared within 30 min after the 4^th^ VI training. The VR+VR group was introduced as a control matched for total training days. AMPA/NMDA ratios were recorded from the CS-projecting LOFC layer 5 neurons **(D)** and BLA-projecting LOFC layer 5 neurons **(E)**. Black and blue dashed lines show the mean value of the Ctrl and VR groups in (B,C) respectively. (D) N=9–10; (E) N=9 for each group. **(F,G)** Correlation between the execution index and AMPA/NMDA ratios in the VR+VI_High_ and VR+VI_Low_ groups. N=18–19. **(H)** Schematic diagram of the two-step training protocol with CALI. Mice received CALI (illumination with 1–2 mW amber light for 1 min) within 5-min after each VI training (day 10–13). **(I)** Representative image of CFL-SN expression and optic fiber implantation in the LOFC. The retroAAV-CaMKII-CFL-SN or retroAAV-CaMKII-SN was injected into the CS and optic fibers were implanted into the LOFC. **(J)** Execution index during VI training with CALI. **(K)** Execution index on day 13 and the fraction of VR+VI_High_ and VR+VI_Low_ mice in each group. SN, N=26; CFL-SN, N=29. **(L)** Effects of CALI in the CS-projecting LOFC neurons on lever pressing in the devaluation test. SN, N=10; CFL-SN, N=10. **(M)** Within-subject changes in the devaluation index of (L). *P < 0.05; **P < 0.01; N.S., not significant.

### Training-induced persistent synaptic potentiation of the CS-projecting LOFC neurons maintains the execution of habitual behavior

To further investigate the causal relationship between the potentiation of CS-projecting LOFC neurons and action execution during habitual behavior, we expressed CFL-SN in the CS-projecting LOFC neurons and erased LTP in these neurons after each VI session (days 10–13; Fig. 6H & I). After this period, the average execution index and proportion of VR+VI_High_ mice in the CFL-SN group was significantly lower than the control (SN) group (Fig. 6J & K). Additionally, the effects of LTP erasure by CALI on the expression of habitual behavior was assessed by devaluation and omission tests. In the devaluation test, there was no significant difference in the number of lever presses between the valued and devalued conditions in both the SN and CFL-SN groups (Fig. 6L). This resulted in the comparable devaluation indices between the SN and CFL-SN groups (Fig. 6M). In addition, CALI in CS-projecting LOFC neurons had no significant effect on motivation toward sucrose reward (Fig. S6A & B). Likewise, in the omission test (Fig. S6C & D), CALI had no significant effect on the decay of lever press rate quantified as reduction in relative lever pressing rate (Fig. S6C & D).

Next, to test if the increased excitability of CS-projecting LOFC neurons during habitual task performance is sufficient for facilitation of action execution, we performed chemogenetic activation of CS-projecting LOFC neurons during VI session (Fig. S6E–L). After completion of the 13-day VR+VI task, mice received additional two VI sessions with CNO or vehicle injection. CNO administration induced a slight but significant increase in lever pressing frequency (Fig. S6F), without affecting frequency of reward acquisitions or nose pokes (Fig. S6G & H). The same manipulation of CS-projecting LOFC neurons had no significant effect on lever-press responses or motivation toward sucrose reward in devaluation test (Fig. S6I–L).

These results indicate that training-induced potentiation in CS-projecting LOFC neurons during habit formation, specifically determines changes in execution level without affecting the transition from goal-directed to habitual strategies.

### Reward-related activity in the CS-projecting LOFC neurons after habit formation correlates with the execution index

To confirm this idea, activity of CS-projecting LOFC neurons during training were recorded using Ca^2+^-imaging (Fig. 7A). At each step of the training, significant responses were associated with both lever pressing and rewarded-nose pokes (Fig. 7B–D). Responses to lever pressing and rewarded-nose pokes arose in two non-overlapping populations of neurons, which were classified as lever press and reward neurons respectively (Fig. 7E & F; Methods). This contrasts with RSC-projecting ACC neurons which responded only to the reward, but not lever pressing (Fig. 5A–G). In addition, the reward-responding CS-projecting LOFC neurons showed a strong response with slow decay following the onset of rewarded nose poke (Fig. 7B, C). This also highlights distinct characteristics from reward-excited RSC-projecting ACC neurons, which frequently exhibit multiple, brief activities during reward consumption (Fig. 5B–D). The overall activity of neurons during lever pressing was unchanged throughout the training session (Fig. 7D & E). In contrast, the activity during the rewarded-nose poke increased specifically in the VR+VI_High_ group, but not in the VR+VI_Low_ group (Fig. 7D). These neurons were mutually inhibited; the lever press neurons were inhibited during the firing of reward neurons and *vice versa* (Fig. 7F). Following a transition from goal-directed to habitual behavior, significant changes in the fraction of both classes of neurons were observed (Fig. 7E) but no correlation was observed between the fractions of cells and execution index of each mouse (Fig. S7A, B). Lever press cell activity was relatively stable at different phases of training, with the exception of a small but statistically significant decrease during the VR training (Fig. 7E & G). There was no significant change in the lever press-related normalized AUC of the lever press cells and correlation with execution index of each mouse (Fig. 7H, Fig. S7C). In contrast, the amplitude of the reward-related Ca^2+^ transients progressively increased across the training stages (Fig. 7E & I). Additionally, at the VI training stage, but not VR stage, the reward-related response had a positive correlation with the execution index of each mouse (Fig. 7J), suggestive of a causal relationship between reward-related activity of CS-projecting LOFC cells and execution index of habitual behavior. Comparing the reward-related activity of each mouse across training stages revealed no consistent enhancement between the VR and VI training stages, suggesting that the positive correlation with the execution index cannot be attributed solely to an increase in the number of training sessions (Fig. S7D). Taken together with the whole-cell recording and LTP erasure data (Fig. 6), reward-related activation of CS-projecting LOFC neurons plays a pivotal role in sustaining a consistent level of execution during habit formation.

**Figure 7.**
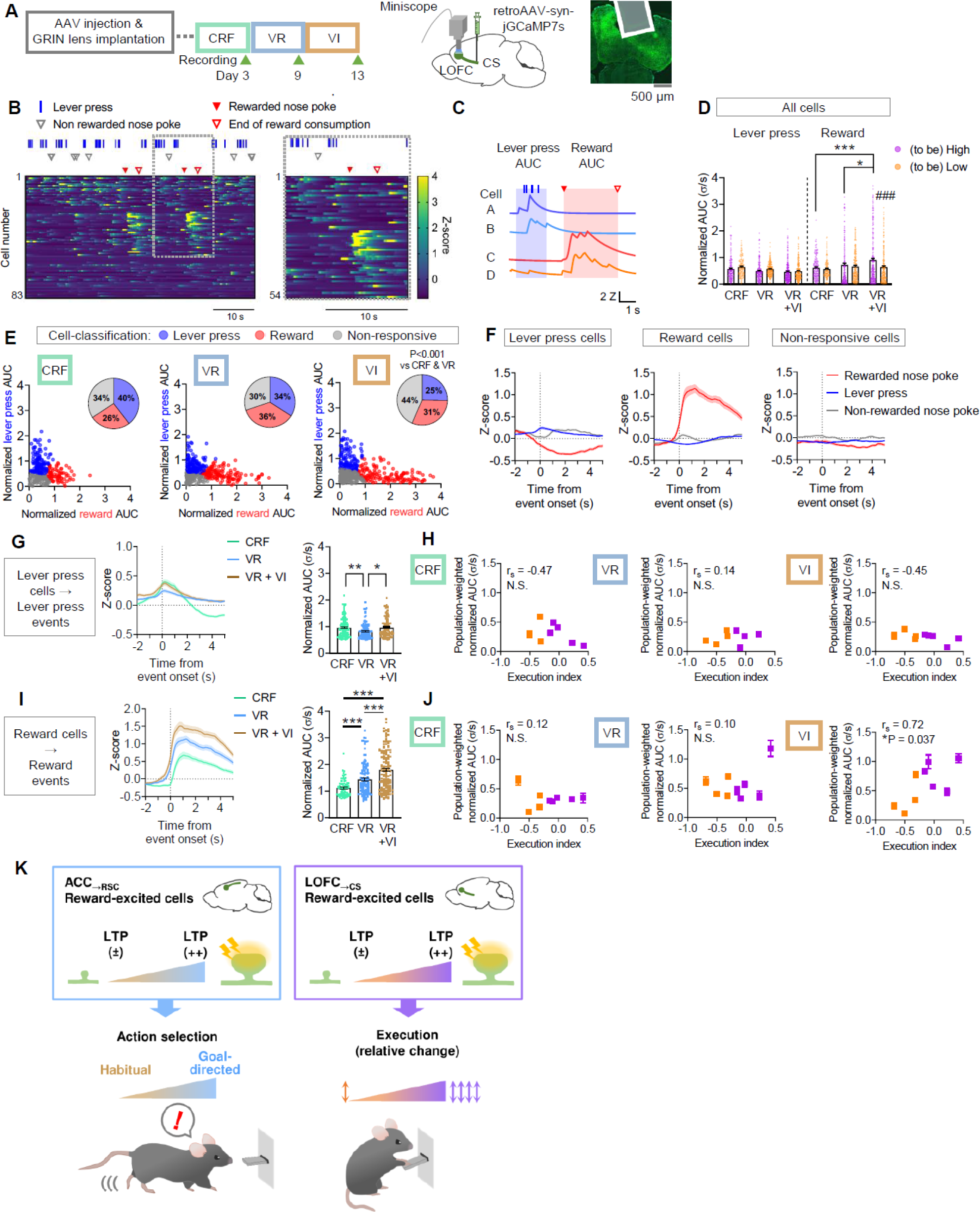
Reward-related activity of CS-projecting LOFC neurons is positively correlated with performance changes during habit formation. **(A)** Schematic diagram of *in vivo* Ca^2+^ recording and representative image of jGCaMP7s expression and GRIN lens implantation. The retroAAV-syn-jGCaMP7s was injected into the CS and GRIN lens was implanted into the LOFC. Recordings were performed in the last CRF (day 3), VR (day 9) and VI (day 13) training sessions. **(B)** Representative heat map of Ca^2+^ transients during VR training (day 9). The right panel shows the enlarged version of the dotted area on the left. **(C)** Representative trace of Ca^2+^ transients around lever pressing and reward events. Cells A and B were activated during lever presses, but not at reward consumption. In contrast, cells C and D showed higher activity during reward consumption than at lever presses. The red and blue highlighted zones show the time windows for calculation of reward and lever press-related neural responses, respectively (see Methods). **(D)** The AUC of event-related responses were normalized by the duration of lever press bouts, or reward consumption. Each dot represents each cell’s average of normalized AUCs for lever press or reward events during training. CRF (to be High), N=166; CRF (to be Low), N=140; VR (to be High), N=179; VR (to be Low), N=208; VR+VI_High_, N=247; VR+VI_Low_, N=213. **(E)** The XY-planes represent lever press and reward-related normalized AUCs for each cell. Cells were classified into three subgroups according to the magnitude and preference of their responses toward lever press and reward events (see Methods). The pie charts represent fractions of each subgroup. **(F)** Average activity of lever press cells, reward cells and non-responsive cells during each type of event during VR training. **(G)** Average temporal dynamics and normalized AUCs of lever press cells to lever press events during each training stage. CRF, N=121; VR, N=132; VR+VI, N=116. **(H)** Correlation between the execution index and population-weighted AUCs of the lever press cells to lever press events during each training stage. CRF, N=121 cells/8 mice; VR, N=132 cells/8 mice; VR+VI, N=116 cells/9 mice. **(I)** Average temporal dynamics and normalized AUCs of the reward cells to reward events during each training stage. CRF, N=80; VR, N=138; VR+VI, N=144. **(J)** Correlation between the execution index and population-weighted AUCs of the reward cells to reward events during each training stage. CRF, N=80 cells/9 mice; VR, N=138 cells/9 mice; VR+VI, N=144 cells/9 mice. *P < 0.05; **P < 0.01; ***P < 0.001; N.S., not significant. For (D), ^#^P < 0.05 at (to be) High vs. (to be) Low. **(K)** Dynamic regulation of cortical region-specific synaptic potentiation orthogonally controls habitual behavior. LTP in RSC-projecting ACC neurons is necessary for the predominance of goal-directed system for action selection, and as it weakens, behavior becomes habitual. By contrast, LTP in CS-projecting LOFC neurons serves to maintain the execution of the habitual behavior at a comparable level to its previous, goal-directed execution.

## Discussion

Identifying the regulatory mechanisms that operate during the transition from goal-directed to habitual behavior has proven difficult, primarily due to the lack of an appropriate behavior model where habit formation can be induced at a defined time window. Our newly designed task, which sequentially combines VR and VI tasks, overcomes this limitation by including clearly separable goal-directed and habit formation phases. Using this task, we demonstrated that habitual behavior should be considered from two mechanistic dimensions, the decision-making strategy and execution level, that are independently controlled. Supporting this, further analyses revealed that decision-making strategy and execution level are orthogonally represented in the brain. At the circuit level, the emergence of goal-directed behavior is mediated by the potentiation of synaptic transmission in RSC-projecting ACC neurons. Then the transition from a goal-directed to habitual strategy is accompanied by a depotentiation of transmission in those same neurons. In contrast, execution level is determined by the persistence/absence of synaptic potentiation in CS-projecting LOFC neurons. We confirmed the necessity and sufficiency of these circuits by chemogenetically manipulating the activity, and additionally by preventing training associated plastic changes. Furthermore, *in vivo* Ca^2+^-imaging revealed changes in neuronal firing patterns, which together demonstrate that distinct populations of prefrontal cortical neurons govern predominance among decision-making strategies and execution level of the habitual behaviors independently.

The ACC is required for monitoring and prediction of an action-outcome relationship^30,31^, by representing various reward-related information, such as reward value and cost of action^32–34^. Consistent with this, we found that chemogenetic manipulation of ACC activity bidirectionally regulated goal-directed/habitual decision strategy. Though recent studies have demonstrated that striatum-projecting ACC neurons are essential for cost-benefit assessment, but not goal-directed decision-making^35^, and BLA-projecting neurons represent the innate value of reward and aversive signals^36,37^, the mechanisms governing goal-directed decision-making have remained elusive. In this regard, this is the first demonstration that plasticity in RSC-projecting ACC neurons can dictate goal-directed strategies for decision-making. Since both the ACC and RSC are crucial for learning and memory of associative information, which is transferred from the hippocampus^28,38^, it is conceivable that information of action-outcome association for goal-directed decision-making, which is transiently encoded in the hippocampus at earlier stages of training^39^, is transferred to the RSC-projecting ACC neurons.

Erasure of the VR training-induced LTP in the RSC-projecting ACC neurons did not impair the acquisition of lever pressing behavior, while facilitating habit formation. Consistent with this observation, previous behavioral studies in rodents have consistently shown that disruption of brain regions implicated in goal-directed strategy, including the dorsomedial striatum, preserves reinforcement learning, yet facilitates habit formation^5,40^. These findings suggest that when the goal-directed strategy is compromised, animals instead recruit reinforcement learning processes biased toward habitual behavior. Supporting this, computational studies have proposed two parallel forms of reinforcement learning, model-based and model-free (or value-free) learning corresponding to goal-directed and habitual strategies respectively, that develop concurrently during behavior acquisition^2,41,42^. Moreover, human studies have shown that habit formation is blocked while model-based learning is predominant, suggesting inhibitory mechanisms of the goal-directed strategy on the habitual strategy^43^. Accordingly, the intact reinforcement learning in the LTP erasure group is most consistent with preferential disruption of model-based learning system.

In contrast to the ACC, although a number of studies implicate the OFC in reward-related decision-making, its functional significance in goal-directed decision-making has been controversial; both activation and inhibition of the OFC disrupt flexibility of a behavior and shift it to a habitual one^5,44^. In addition, other prior studies have shown the OFC has no, or a limited contribution to behavior flexibility and defining decision strategies^45,46^. While possible causes for this inconsistency, such as heterogeneity of the OFC subdivisions and testing protocols, have been postulated^5,44,45,46^, a specific role for the OFC in goal-directed strategies is still to be fully elucidated.

Despite the functional differences between RSC-projecting ACC neurons and CS-projecting LOFC neurons, Ca^2+^-imaging revealed that both neurons represent reward (outcome)-related information. Considering that *post hoc* LTP erasure was sufficient to affect the decision strategy and execution level, training-induced learning and updating of outcome-related association, rather than innate representation of reward value, may be critical for the effects of both neurons. As conceptualized by computational studies^2,41,46^, rewarded experience updates the model for stimulus-response-outcome associations when using a model-based learning strategy. Hence, outcome-related information encoded in the RSC-projecting ACC neurons is likely to be utilized for updating the model of stimulus-response-outcome associations.

In contrast, outcome-related information encoded in the CS-projecting LOFC neurons is less likely to directly modulate the habitual strategy, since this system utilizes stimulus-response associations rather than action-outcome associations. In line with this, comparative studies of ACC and OFC highlight their dissociative roles in encoding outcome value; the OFC encodes outcome-related information during Pavlovian (cue-outcome) learning rather than during the instrumental (action-outcome) learning, while the opposite contribution is found in the ACC^47,48^. Pavlovian learning is known to facilitates execution of operant behavior through Pavlovian-instrumental transfer, particularly when the response is habitual^49,50^. Although our task was not explicitly designed to induce Pavlovian conditioning, interoceptive cues during tasks such as hunger could be associated with reward presentation. This raises the possibility that cue-outcome encoding in the CS-projecting LOFC neurons facilitates execution level of habitual behavior. In addition, it has been reported that the OFC is critical for contextual knowledge of outcome expectation, which hierarchically regulates performance of already acquired behaviors^51,52^. This function of the OFC has been mainly discussed in relation to flexible adaptation of behaviors according to context discrimination, where subjects learn distinct associations in a context-specific manner^5,52^. By contrast, when a behavior is stably rewarded in a specific context, this contextual stability has been proposed to facilitate execution of a habitual behavior, possibly through serving the contextual information as a promoter^8,14,53^. Taken together, LTP in CS-projecting LOFC neurons could contribute to the association between rewarded experience and its preceding cue or context, thereby facilitating the execution of lever pressing habit without affecting the stimulus-response associations.

The regulation of goal-directed and habitual behaviors by cortico-striatal pathways is well established^8,10^. Accumulating evidence supports dichotomous control mechanisms, such as “dorsomedial versus dorsolateral striatum” and “associative versus sensorimotor circuits”^54,55^. However, recent histological analyses revealed distinct projections to striatal regions from each cortical region^56,67^ suggesting that these histologically separated pathways have distinct functions. Consistently, it has repeatedly been reported that LOFC neurons send inputs to a subregion of the striatum ranging from the dorsomedial to the central striatum. Dense inputs are observed in the central subregion^57–60^. This unique projection pattern of the LOFC supports the present finding that CS-projecting LOFC neurons are implicated in habit execution rather than decision-making strategy.

Accumulating evidence from animal and clinical research has shown that hyperactivity in the OFC-striatum pathway plays a pivotal role in the pathology of compulsivity^61–63^. Abnormally strong habit is considered as a possible cause of excessively repeated execution of certain behavior(s) shown in patients with compulsivity^64,65^. Symptom-related, cue-induced hyperactivity in the OFC and increased functional connectivity between the OFC and striatum have been reported in patients with compulsive disorders^66–69^. While present study focuses on habitual behavior in healthy mice, the concept of LOFC-derived individuality in habit execution may be applicable to the pathological habitual behaviors such as obsessive-compulsive disorder, addiction and eating disorders.

In conclusion, the present study provides novel insights into neural mechanisms governing behavioral control during the goal-directed to habitual transition. The spatiotemporally distinct online encoding of outcome-related information in the ACC and LOFC was required for blocking habit formation and maintenance of action execution, respectively (Fig. 7K). Our findings highlight the ACC-RSC pathway as a critical determinant of decision-making strategy, and further, reveal an unexpected role of LOFC-CS pathway on controlling habit execution.

## Acknowledgments

We thank Drs. Dai Watanabe, Tomoyuki Furuyashiki, Chika Nishimura, and Steven Middleton for their comments on the manuscript. This work was supported in part by the Japan Society for the Promotion of Science Grants-in-Aid for Scientific Research JP22K15291 and JP25K02556 from the MEXT, Japan, AMED Brain/MINDS 2.0 JP24wm0625507, Uehara Memorial Foundation, Kobayashi Foundation, Takeda Science Foundation and Astellas Foundation for Research on Metabolic Disorders (to N.A.) and Grants-in-Aid for Scientific Research JP18H05434, JP22H04981 and JP22K21353 from the MEXT, Japan, The Uehara Memorial Foundation, The Naito Foundation, Research Foundation for Opto-Science and Technology, Novartis Foundation, and The Takeda Science Foundation, HFSP Research Grant RGP0022/2013, and JST CREST JPMJCR20E4 (to Y.H.).

## Author contributions

N.A. and Y.H. designed the research. N.A. and D.P. performed the experiments and collected data. N.A. and Y.H. conceived and conducted behavioral and neural analyses. N.A. and Y.H. wrote the manuscript.

## Supplemental information

Document S1. Figure S1–7.

Table S1. Excel file containing statistical information.

## Materials and Methods

### Animals

All animal care and experimental procedures were conducted in accordance with the ethical guidelines of the Kyoto University Animal Research Committee. Male C57BL/6J mice (RRID: IMSR_JAX:000664) were purchased from Japan SLC (Shizuoka, Japan). Mice (6–24 weeks old) were housed at a constant ambient temperature of 24 ± 1°C on a 12-h light-dark cycle with access to food and water *ad libitum*. When food restriction was conducted, food intake was limited to 2–2.2 g/day on weekdays (80–90% of the *ad libitum*-fed body weight). All behavioral experiments were randomized and videotaped.

### Operant training

Operant training was carried out using an operant chamber (Med Associates, St Albans, VT) controlled by MED-PC software (version IV and V; Med Associates). Each session began with turning on the house light and lever presentation and ended when mice received the maximum number of reinforcers, or when 60 min had elapsed, unless otherwise described. For magazine training (2 or 3 (for Ca^2+^ recordings) sessions), food-restricted mice were placed in an operant chamber without lever presentation, after which 16 reinforcers (10 μL of 20% sucrose solution) were presented at 60-s intervals on average. For habituation to the lever, mice received three continuous reinforcement (CRF) sessions, where each lever press was rewarded. The max number of reinforcers increased across sessions (5, 15 and 30 reinforcers). Following CRF sessions, mice were trained with the VR or VI protocols. During VR training, an average of 10 (VR10) or 20 (VR20) presses were required for each reinforcer, whereas for VI training the reward was delivered on first press after an average interval time of either 30 (VI30) or 60 (VI60) sec following delivery of the previous reward. The sets of variables were fixed, but their order was changed across sessions. Mice that underwent the VR10 or VI30 schedule for the first two VR or VI sessions; thereafter were trained with the VR20 or VI60 schedule. When mice were unable to obtain 10 reinforcers on the 4^th^ VR20 or VI60 session, they were excluded from further experiments.

For the two-step paradigm, mice received three CRF, two VR10 and four VR20 sessions, which were followed by four VI60 sessions. The transition from VR20 to VI60 schedules were seamlessly carried out; without any cue or context changes. The changes in lever press rates between the last VR (fourth VR20) session and last VI60 session (fourth VI60 session) were indexed as follows; Execution index = [(press rate in VI session) – (press rate in the VR session)]/ [(press rate in the VI session) + (press rate in the VR session)].

For Fig. 7C, press rate in each VI60 session, was used instead of the last VI60 session.

All lever presses, reward deliveries and nose pokes during training were automatically recorded by the MED-PC software. The end of reward consumption was manually detected as the time at which mice start to move their head away from the reward port. For analyzing lever press bouts, presses with an inter-event interval of < 0.5 s were considered as one bout.

### Devaluation test

After the last training session, devaluation tests were conducted over two consecutive days. Mice were allowed free access to food chew (the valued day) or a 20% sucrose solution (the devalued day) for 30 min. The order of the valued and devalued days was randomized. After the free access session, mice were placed in the operant chamber for 5 min where the lever was presented as usual, but never reinforced. To normalize the baseline variability of lever pressing activity among individuals, the difference in lever pressing numbers between the valued and devalued conditions for each mouse was presented as the devaluation index. The devaluation index was calculated as follows:

Devaluation index = [(lever presses on the valued day) – (lever presses on the devalued day)]/ [(lever presses on the valued day) + (lever presses on the devalued day)].

One mouse was excluded from experiments shown in Fig. S6J–M because the pellet intake during the free access session was too low to detect.

### Omission test

After the last training session, an omission test was conducted. During the 30-min omission session the lever was presented as usual. When the lever was not pressed for 20 s, a reward (20% sucrose solution) was provided at the reward port, however when the lever was pressed a 20-second cooldown timer was reset.

For analysis, press rate was calculated every 5 min and divided by press rate in the last VI60 session. The time to reach 50% and 75% of the total press number was used to compare the decay of lever pressing activity.

### Spatial discrimination learning and reversal learning

After completion of the two-step operant training, T-maze spatial discrimination training was performed as previously described^70^. Food restriction was continued throughout all behavioral experiments. Mice were habituated with the T-maze apparatus (one start arm (30 × 10 cm), two goal arms (30 × 10 cm) and 30-cm-high surrounding walls) for 7 days and trained for spatial discrimination for 12 days (5 trials/day). The rewarded arm (rewarded with 100 μL of sweetened milk) was randomly chosen and fixed during the 12-day training sessions. For reversal learning, the rewarded arm was reversed, and mice underwent 10 free-choice trials per day for 4 days.

### Marble burying test

After completion of the T-maze experiments, compulsive-like behavior was tested by the marble burying test. Each mouse was introduced to a novel plastic cage with 5-cm cage bedding (wood chips) and 15 glass marbles. The number of marbles whose surface were covered more than 50% with cage bedding were counted across the 10-min session.

### Stereotaxic surgery

Mice were anesthetized with isoflurane (1–2%) and fixed on a stereotaxic frame (Narishige, Tokyo, Japan). Adeno-associated virus (AAV) vector (0.3 μL/side) was injected with a stainless-steel cannula (30 G) at the following coordinates, unless otherwise described; ACC (AP 1.2 mm, ML ±0.5 mm, DV 1.7 mm), LOFC (AP 2.7 mm, ML ±1.7 mm, DV 2.5 mm), RSC (AP -1.0 mm, ML ±0.4 mm, DV 0.8 mm), CS (AP 1.2 mm, ML ±1.6 mm, DV 4.0 mm) and BLA (AP -1.5 mm, ML ±3.2 mm, DV 4.5 mm). The injection site for the CS was selected based on previous reports^58–60^.

### DREADD experiments

For global manipulation of ACC/LOFC pyramidal neurons, the AAV vector expressing hM4Di (AAV-CaMKIIα-hM4Di-mCherry; Addgene #50477) or hM3Dq (AAV-CaMKIIα-hM3Dq-mCherry; Addgene #50476) was bilaterally injected into the ACC or LOFC. For manipulation of the CS-projecting LOFC neurons, the AAV vector expressing hM3Dq (AAV-DIO-hSyn-hM3Dq-mCherry; Addgene #44361) and retrograde AAV vector expressing Cre (AAV.CamKII 0.4.Cre.SV40; Addgene #105558) was bilaterally injected into the LOFC and CS, respectively. Mice were allowed to recover for at least one week prior to starting the operant training; so that at least 4-weeks had passed before CNO was injected.

For the devaluation tests, CNO (0.5 mg/kg; Cayman Chemical, MI, USA) or its vehicle (1% DMSO in saline) was injected prior to starting the free access session. The dose of CNO was chosen based on previous studies^5, 72^. For the cross-over design, mice were re-trained with one VR20 session after the first set of devaluation tests. The second set of devaluation tests were performed under the opposite drug condition. The order of the valued and devalued conditions was the same as in the first set. For manipulation during the training and omission test, CNO or its vehicle were injected 20 min before starting the session.

### In vivo Ca^2+^ imaging

Mice received unilateral injections of retroAAV-syn-jGCaMP7s-WPRE (Addgene #104487) in either the RSC or CS. To obtain a sufficient number of jGCaMP7s-positive cells, AAV was injected at two sites within the RSC or CS; RSC (AP -1.0 & -2.0 mm, ML ±0.4 mm, DV 0.8 mm) and CS (AP 1.2 & 0.2 mm, ML ±1.7 mm, DV 4.0 mm). Next a GRIN lens (1-mm diameter, 4-mm length; Inscopix, CA, USA) was placed in the ACC (AP 1.2 mm, ML ±0.8 mm, DV 0.5 mm) or LOFC (AP 2.7 mm, ML ±1.7 mm, DV 1.0 mm), after which mice were singly housed until experiments concluded.

At least 3 weeks post injection, a baseplate for the UCLA Miniscope^29^ (version 4.4 purchased from OpenEphys) was fixed over the GRIN lens with dental cement. Lens implantation and baseplate mounting were performed with slight modifications to the method described at http://Miniscope.org. Operant training began at least 4 weeks after AAV injection. To enable mice with a Miniscope to enter the reward port, a wide custom-made nose poke hole was used. Mice underwent the two-step operant task with three magazine training sessions. For habituation to the Miniscope, mice were trained with a dummy scope after the second magazine training session. On the 3^rd^ CRF, 6^th^ VR and 4^th^ VI sessions (the last session of each training schedule), calcium transients during the session were recorded at 30 Hz using the UCLA Miniscope DAQ software (version 1.10). Recordings of calcium transients and animal behaviors were started and finished simultaneously with the operant training sessions, by TTL signals from the MED-PC V software. Contrast and brightness of the videos were adjusted using FFmpeg and semi-automatically processed with the MiniAn pipeline^71^ for removing sensor noise and background fluorescence, motion correction and source separation were performed by a constrained non-negative matrix factorization (CNMF) algorithm. The spatial footprint and calcium signal for each detected cell were manually verified to exclude false-positive detections. While we maintained the same imaging fields across sessions, individual neurons were not longitudinally tracked due to minor mechanical displacements of the imaging window and the focal plane between recording days. Instead, all comparisons across training stages were conducted as population-level analyses.

### Event-related Ca^2+^ activities

Following the MiniAn pipeline, output values of Ca^2+^ activity were normalized by their standard deviation (σ). For comparisons of the average Ca^2+^ responses (Fig. 5E, G, H & Fig. 7G, H, J), activity was Z-scored to highlight event-related activity changes from mean activity. Z-scores were calculated as follows; Z-score = (C – μ) / σ, where C is an output value from the MiniAn pipeline and μ is mean of the overall output values. The time windows for lever pressing, non-rewarded nose pokes and rewarded nose poke events were determined as follows; Lever press: A cluster of lever presses (inter-event interval (s) < 0.5 s) was treated as a single bout. The time window for analysis was set from 0.5 s before a bout, to 0.5 s after a bout. When mice nose poked during this window, the event was excluded.

Rewarded nose poke: The first nose poke after reward delivery was considered as a rewarded nose poke. The time window from detection of rewarded nose pokes, to end of the reward consumption (manually detected as described above) was used for analysis. In the LOFC recording group, nine events (2 mice, CRF session) were excluded because the sensor detected objects other than mouse’s nose (including the Miniscope, paw and tail). The pre-event period was set at 0.5 seconds prior to an event. Since the length of each event varied significantly, even within a single recording session, event-related Ca^2+^ activity was normalized by event length (time-normalized AUC). For excitation, the mean Ca^2+^ activity during events (σ) was divided by event length. For inhibition, event-related negative AUC below the mean Ca^2+^ activities of pre-event periods, was divided by event length.

Non-rewarded nose poke: To avoid contamination of rewarded nose poke events, nose pokes that performed between lever presses that did not trigger reward presentation were classified as a non-rewarded nose poke. Based on this definition, non-rewarded nose pokes were analyzed only in the VR and VI sessions. The time window was set to ±0.5 s around the nose poke, if further nose pokes occurred within a 0.5 s interval, only the first one was included in analyses.

### Analysis of Ca^2+^ transients in RSC-projecting ACC neurons

RSC-projecting ACC neurons were separated into three subclasses (reward-excited, reward-inhibited and non-responsive cells).

Reward-excited cells: Neurons which met the following three criteria were classified as reward-excited cells; (1) > 1 σ peak amplitude in at least 20% of rewarded nose poke events, (2) mean amplitude during rewarded nose pokes was larger than that during pre-nose poke periods, (3) showing multiple peaks during at least one rewarded nose poke event (Fig. 5D). Since, typical rewarded nose pokes lasted 2-20 s, the criteria (2) and (3) were used to exclude a subset of neurons which showed non-event-related episodic Ca^2+^ activity.

Reward-inhibited cells: Neurons that were not classified as reward-excited cells were further tested for their decreases in Ca^2+^ activities during rewarded nose poke events. For each cell, the minimum value of Ca^2+^ activity during each rewarded nose poke event was compared with the mean amplitude during the corresponding pre-nose poke period using a paired *t*-test. When there was a significant decrease in Ca^2+^ activity, the neuron was classified as a reward-inhibited cell.

For further statistical analyses, excitatory/inhibitory magnitude of each reward-excited/inhibited cell was scored as a population-weighted response (Fig. 5I & J), which was calculated by normalized AUC of rewarded nose pokes, multiplied by the fraction of reward-excited/inhibited cells in each mouse^73^.

### Classification of activity of CS-projecting LOFC neurons

CS-projecting LOFC neurons were separated into three subclasses (reward, lever press and non-responsive cells), as follows.

First, normalized AUCs of Ca^2+^ activities for reward and lever press events were calculated. Among neurons recorded in the VR session, time-normalized AUCs for each event were Z-scored, so that features of response selectivity could be compared. To this end, neurons recorded in the VR session were clustered into 5 clusters by the K-means+ method (Python with scikit-learn package). The number of clusters was determined by the elbow method. Then, to apply this classification to recordings in the CRF and VI sessions, we made a logistic regression model for classification of neurons using time-normalized AUCs datasets. For training, 75% of the datasets in the VR session were used. The accuracy of the model tested with the remaining 25% of the datasets was 0.89.

Using the classification, neurons were separated into 5 groups: with high and moderate preference for lever pressing, high and moderate preference for reward and non-responsive for both events. Since the numbers of high preference neurons were small, high and moderate preference groups for each event were combined. For each event-related neurons normalized AUC and population weighted normalized AUC were calculated as described above.

### CALI experiments

pAAV-CaMKIIα-CFL-SN and pAAV-CaMKIIα-SN were constructed by replacement of the hM4Di-mCherry gene in the pAAV-CaMKIIα-hM4Di-mCherry (Addgene #50477) virus, to the CFL-SN and SN genes, respectively. Retrograde AAV vectors for CFL-SN and SN expression were produced as previously described^70^.

Mice received bilateral injections of retroAAV-CaMKIIα-CFL-SN or retroAAV-CaMKIIα -SN in the RSC or CS. Fiber optic cannulae, constructed from ceramic ferrules (1.25 mm diameter; CFLC270-10; Thorlabs NJ, USA) and plastic optic fibers (250 μm diameter; CK-10; Mitsubishi Rayon, Tokyo, Japan), were implanted in the ACC (AP 1.2 mm, ML ±0.5 mm, DV 1.0 mm, angled 10° relative to the ML axis) or LOFC (AP 2.7 mm, ML ±1.7 mm, DV 2.5 mm). 595 nm laser light intensity from the tip of the fiber optic cannula was adjusted to 1–2 mW. Operant training commenced at least 1-week post-surgery, so that at least 3 weeks had elapsed since injection of virus.

For CALI in the RSC-projecting ACC neurons, mice received bilateral illumination of the ACC for 1 min, within 5 min after VR training had ended. For CALI in the CS-projecting LOFC neurons, mice received bilateral illumination of the LOFC for 1 min, within 5 min after VI training had ended.

### Histology

After completing all behavioral experiments, mice were deeply anesthetized with isoflurane and transcardially perfused with PBS followed by 4% paraformaldehyde. Whole brain samples were further fixed in 4% paraformaldehyde for 30 min at 4°C and incubated in 15% sucrose/PBS solution overnight at 4°C. Coronal brain sections (30-μm thick) were prepared with a cryostat (Leica CM1950/Leica CM1860; Leica Biosystems, Nußloch, Germany). Signals from fluorescent proteins were visualized by a confocal microscopy system (FLUOVIEW FV1200, Olympus, Tokyo, Japan; ECLIPSE Ti2-E, Nikon, Tokyo, Japan). For visualizing jGCaMP7s, sections were stained with rabbit anti-GFP antibody (1:500; A11122, lot# 2659306; Thermo Fisher Scientific, MA, USA) followed by AlexaFluor 594 conjugated secondary antibodies (1:200; Cell Signaling Technology, MA, USA). Animals were excluded if post hoc histological analysis revealed viral expression or fiber or lens placement outside the target region.

### Electrophysiology recordings

Electrophysiology recordings were performed as previously described^70,74^ with minor modifications. For the control group, mice were food restricted for at least 2 weeks, but did not receive operant training. In the projection-specific subpopulation groups, mice were injected with retroAAV-CAG-tdTomato (Addgene #59462) at least 3 weeks before recording.

Within 30-min after the last operant training session, mice were deeply anesthetized with isoflurane and decapitated. Coronal brain slices (200-μm thick) were prepared with a vibratome (VT1000S, Leica Biosystems) filled with ice-cold cutting solution (in mM: 120 NMDG-Cl, 2.5 KCl, 26 NaHCO_3_, 1.25 NaH_2_PO_4_, 0.5 CaCl_2_, 7 MgCl_2_, 15 D-glucose, and 1.3 ascorbic acid, pH 7.2). Slices were recovered in ACSF (composition in mM: 124 NaCl, 3 KCl, 26 NaHCO_3_, 1 NaH_2_PO_4_, 2.4 CaCl_2_, 1.2 MgCl_2_, and 10 D-glucose, pH 7.3, oxygenated with 95%O_2_/5% CO_2_) at 32°C for at least 1 h before recording. After recovery, individual slices were transferred to a recording chamber with continuous perfusion of oxygenated ACSF, heated to 27 ± 1°C.

Signals were passed to an EPC9 amplifier (HEKA, Pfalz, Germany) and data were recorded using Patchmaster software (HEKA). Individual neurons were visualized with a microscope equipped with a 40 × water-immersion objective lens (Carl Zeiss, Jena, Germany) and a CCD camera. The resistance of electrodes was 3-7 MΩ when filled with the internal solution (in mM: 120 CsMeSO_4_, 15 CsCl, 8 NaCl, 10 HEPES, 2 Mg-ATP, 0.3 Na-GTP, 0.2 EGTA, 10 TEA-Cl, and 5 QX-314, pH 7.3 adjusted with CsOH). The series resistance was compensated by 70% and maintained at less than 35 MΩ. Additionally, a glass stimulating electrode filled with ACSF was placed near the recording site.

To record EPSCs, membrane potentials were held at -70 mV and a GABA_A_ antagonist (20 μM bicuculline) was continuously perfused. AMPA-mediated evoked EPSCs (eEPSCs) and mixed AMPA and NMDA-mediated eEPSCs were evoked by stimulation at -70 mV and +40 mV, respectively. NMDA-mediated eEPSC amplitude was determined as the average amplitude between 45 and 55 ms post-stimulation. The average of 5 consecutive AMPA- and NMDA-mediated eEPSC measurements was used for analyzing AMPA/NMDA ratios. AMPA receptor (AMPAR)-mediated eEPSCs were normalized by NMDA receptor (NMDAR)-mediated eEPSCs. During paired pulse ratio experiments, electrical stimuli were delivered with a 30-ms interval. The average of 5 consecutive PPRs was used for analysis.

When recording asynchronous EPSCs, CaCl_2_ in the ACSF was substituted with 6 mM SrCl_2_. Asynchronous events evoked by electrical stimulation were recorded 10 times from each neuron and analyzed by Minianalysis software (SynaptoSoft, Decatur, GA) and Win EDR 3.7.3 (University of Strathclyde). Events with amplitudes larger than 15 pA were used for analysis.

### Statistical analyses

Data are presented as the mean ± standard error of mean (S.E.M) unless otherwise stated. Box and whisker plots show the maximum, 75^th^ percentile, median, 25^th^ percentile and minimum values. Statistical analyses were performed with GraphPad Prism 8 (GraphPad, CA, USA). P values < 0.05 were considered significant. All statistical methods, sample sizes and detailed statistics are shown in Supplementary Table 1.

## Data availability

The raw imaging and behavioral datasets are too large to share publicly. However, they are available from the corresponding author upon reasonable request for research purposes. Source data are provided with this paper.

