## Supplemental Figures for "Dissociable roles of prefrontal plasticity in decision making strategy and execution of habitual behavior"

### Supplementary Figures

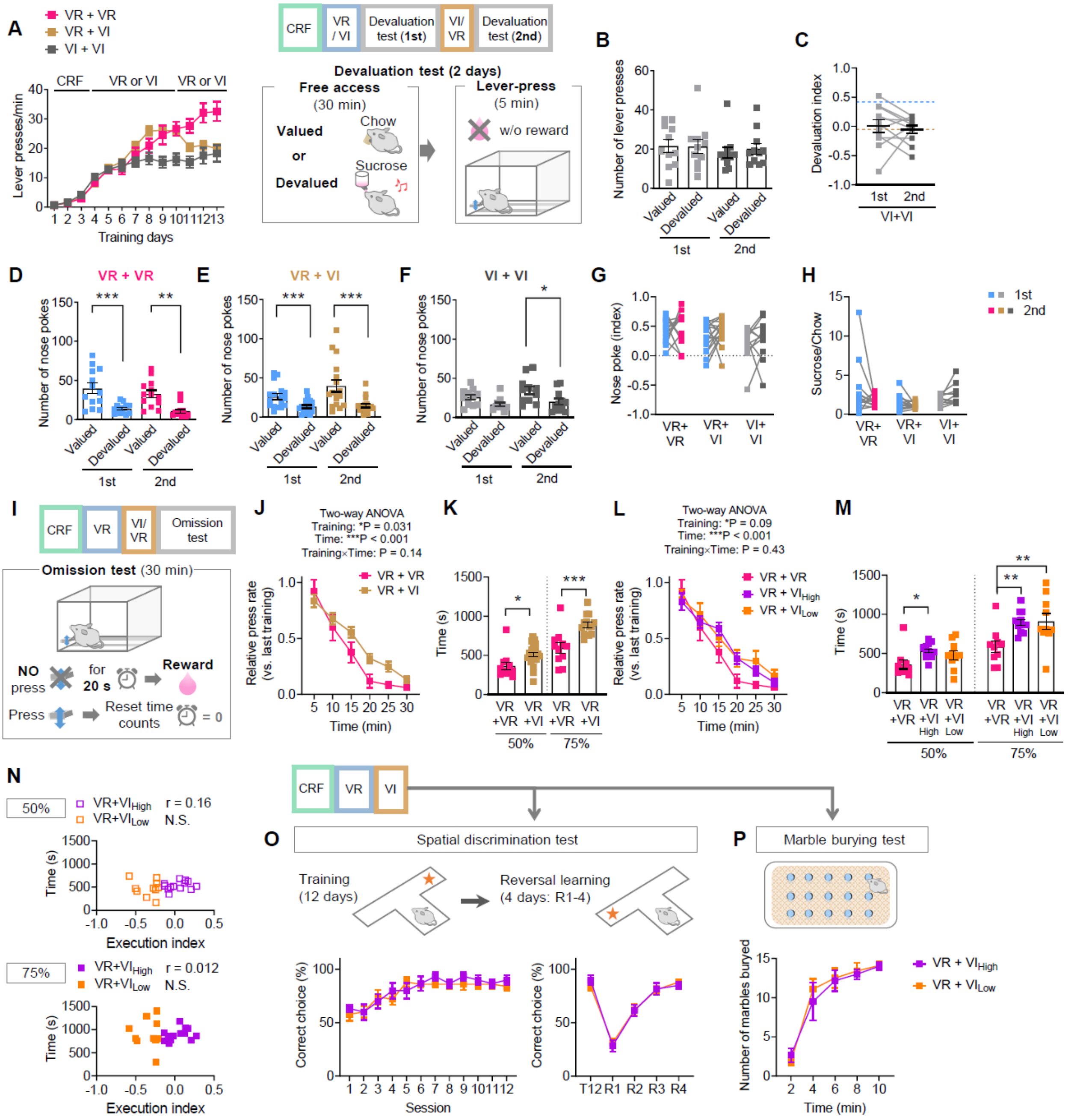


**Figure S1**

*Reward seeking behavior and behavioral flexibility in VR+VR, VR+VI and VI+VI groups, related to Figure 1*

**(A)** For the VI+VI group, mice received 3-day CRF, 2-day VI30 and 4-day VI60 trainings, which were then followed by the first devaluation test. Then, after the additional 4-day VI training, the second devaluation test was performed. Transitions of lever pressing rate in the two-step trainings are plotted with the data from the VR+VR and VR+VI groups, which are same as that shown in Fig. 1B. VR+VR, N= 19; VR+VI, N= 79; VI+VI, N=11. **(B)** Numbers of lever presses on each devaluation test in the VI+VI group. N=11. **(C)** Within-subject differences in lever pressing between valued and devalued conditions, are shown as the devaluation index. The blue and brown dashed lines show the mean values of the 1st and 2nd tests for VR+VI groups in Fig. 1F. N=11. **(D–F)** Number of nose pokes during devaluation tests in mice trained with the VR+VR **(A)**, VR+VI **(B)** and VI+VI **(C)** protocol. (D) N= 13; (E) N= 17; (F) N=11. **(G)** Comparison of the effects of devaluation on nose pokes between the VR+VR, VR+VI and VI+VI groups. Within-subject differences in the number of nose pokes between valued and devalued conditions was indexed as follows; index = [(nose pokes on the valued day) – (nose pokes on the devalued day)/ [(nose pokes on the valued day) + (nose pokes on the devalued day)]. VR+VR, N= 13; VR+VI, N= 17; VI+VI, N=11. **(H)** Ratios of intake of sucrose and chow in the free-access sessions in each group. For (E, G, H), data sets for the VR+VR and VR+VI groups correspond to the experiments shown in Fig. 1D and E. VR+VR, N= 13; VR+VI, N= 17; VI+VI, N=11. **(I)** Schematic diagram of the omission test. Omission test was performed after the 4-day VI training. During the 30-min test session, the sucrose reward was presented every 20 s. Each lever press caused delay of reward delivery. **(J)** Lever press rate during the omission test normalized by the lever press rate in the last training. VR+VR, N= 11; VR+VI, N= 23. **(K)** Time to the 50% and 75% of the total presses during the omission test. VR+VR, N= 11; VR+VI, N= 23. **(L–N)** The data from VR+VI group shown in **(J)** and **(K)** are subdivided into the VR+VI$_{High}$ and VR+VI$_{Low}$ groups according to the execution index. **(L)** Normalized lever press rate. **(M)** Time to the 50% and 75% of the total presses. **(N)** Assessment of the correlation between the execution index and time to the 50% and 75% of the total presses during the omission test. VR+VR, N= 11; VR+VI$_{High}$ N=13; VR+VI$_{Low}$ N=10. **(O)** After the VR+VI trainings, mice were trained with the spatial discrimination task in a T-maze for 12 days. After the 12-day training (5 trials/day), the location of the reward (sweetened milk) was transferred to the opposite arm for the reversal learning test (4 days; 10 trials/day). Among all sessions, there was no significant differences between the VR+VI$_{High}$ and VR+VI$_{Low}$ groups. VR+VI$_{High}$ N=6; VR+VI$_{Low}$ N=10. **(P)** For the marble burying test, mice were introduced into the large cage where 15 glass marbles were placed. The number of marbles which were hidden more than 50% by cage bedding was counted. There was no significant difference between the VR+VI$_{High}$ and VR+VI$_{Low}$ groups. VR+VI$_{High}$ N=6; VR+VI$_{Low}$ N=10. *$P < 0.05$; **$P < 0.01$; ***$P < 0.001$; N.S., not significant.

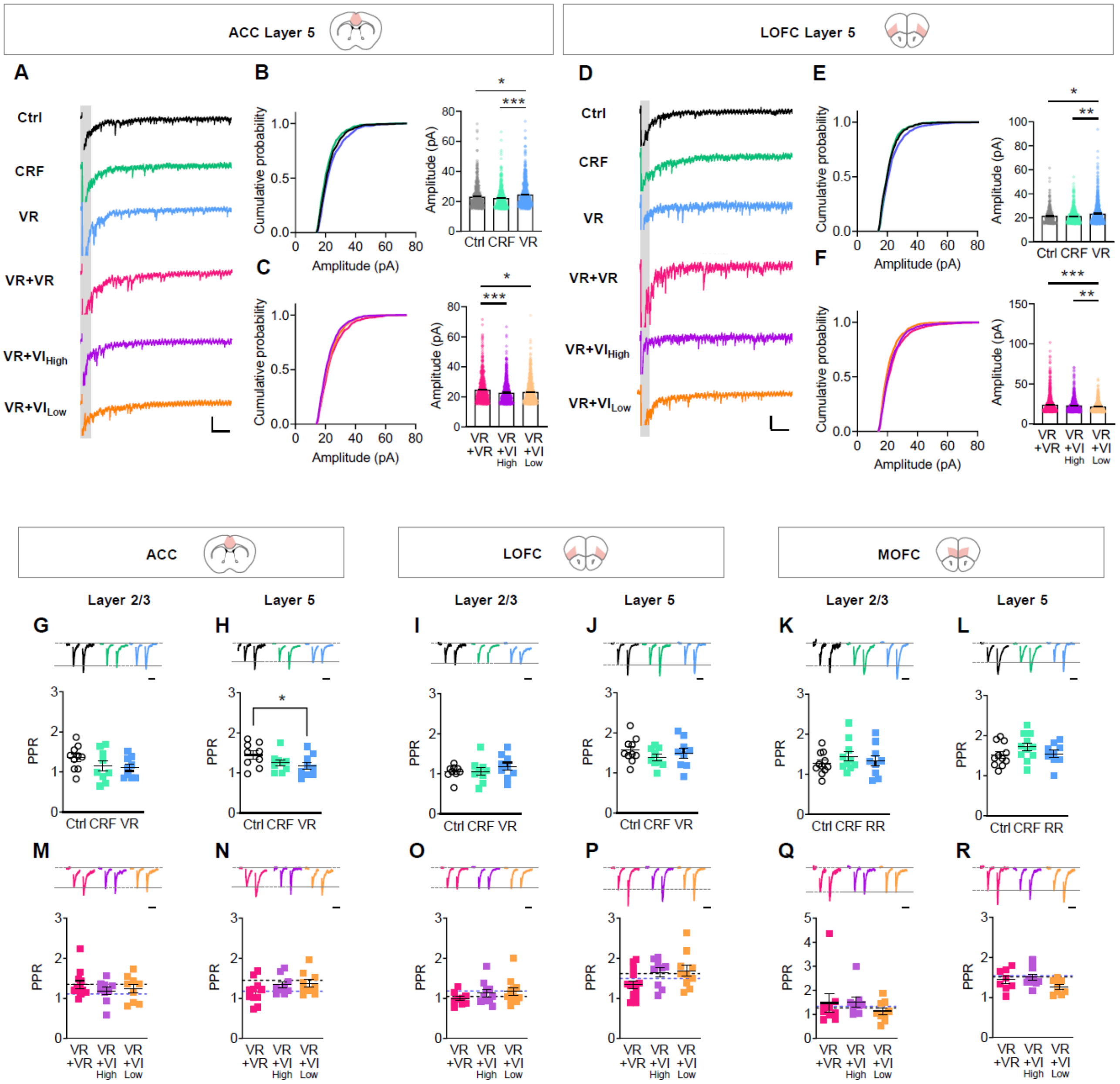


**Figure S2**

*Asynchronous EPSCs and paired pulse ratios in cortical pyramidal neurons, related to Figure 2*

**(A–C)** Acute brain slices of the anterior cingulate cortex (ACC) were prepared within 30 min after the 3rd CRF, 6th VR training **(B)** and 4th VI training **(C)**. Asynchronous EPSCs were recorded from pyramidal neurons in the ACC layer 5. **(D–F)** Acute brain slices of the and lateral orbitofrontal cortex (OFC) were prepared within 30 min after the 3rd CRF, 6th VR training **(E)** and 4th VI training **(F)**. Asynchronous EPSCs were recorded from pyramidal neurons in the LOFC layer 5. The control (Ctrl) mice received food restriction but were not trained. The VR+VR group was introduced as a control matched for total training day. aEPSCs (50 ms-800ms after stimulation) were sampled 10 sweeps per cell. (B) N=447–593 events; (C) N=602–801 events; (E) N=325–709 events; (F) N=632–1215 events for each group. **(G–L)** Acute brain slices of the anterior cingulate cortex (ACC), lateral orbitofrontal cortex (OFC) and medial OFC were prepared within 30 min after the 3rd CRF and 6th VR training. Paired pulse ratios were recorded from pyramidal neurons in the ACC layer 2/3 **(G)**, ACC layer 5 **(H)**, LOFC layer 2/3 **(I)**, LOFC layer 5 **(J)**, medial OFC layer 2/3 **(K)** and medial OFC layer 5 **(L)**. (G) N=9–10; (H) N=9; (I) N=9; (J) N=9–10; (K) N=9–11; (L) N=9–11 for each group. **(M–R)** Acute brain slices of the ACC, LOFC and medial OFC were prepared within 30 min after the 4th VI training. The VR+VR group was introduced as a control matched for total training day. Paired pulse ratios were recorded from pyramidal neurons in the ACC layer 2/3 **(M)**, ACC layer 5 **(N)**, LOFC layer 2/3 **(O)**, LOFC layer 5 **(P)**, medial OFC layer 2/3 **(Q)** and medial OFC layer 5 **(R)**. The black and blue dashed lines show the mean value of the Ctrl and VR groups in **(G–R)** respectively. (M) N=9–12; (N) N=9–11; (O) N=9–11; (P) N=9–14; (Q) N=9; (R) N=8–10 for each group. *P <

0.05; **P < 0.01; ***P < 0.001. Scale bars = 50 pA & 100 ms for **(A & D)**, and 25 ms for **(G–R)**.

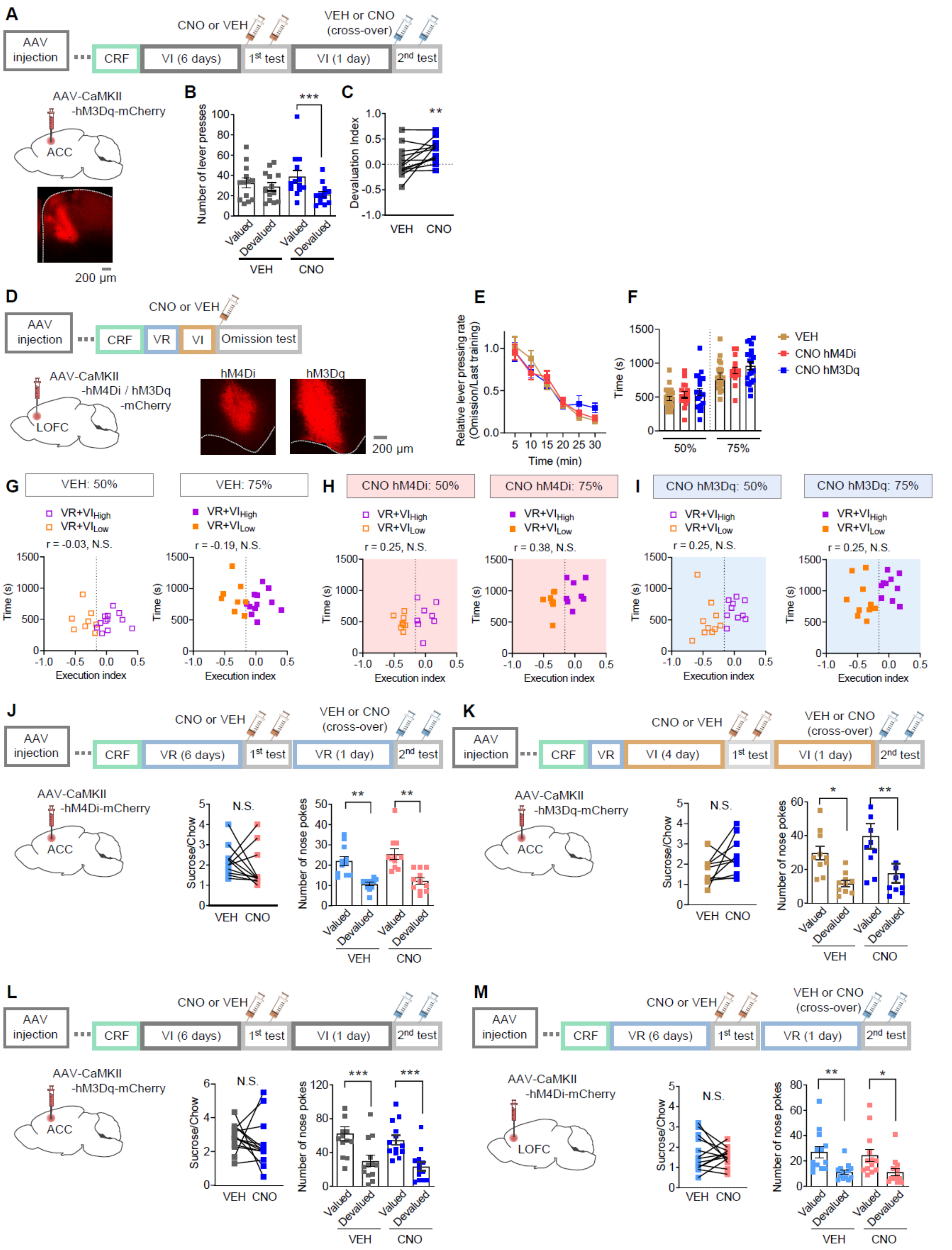


**Figure S3**

*Effects of chemogenetic manipulations of the ACC and LOFC during the devaluation and omission tests, related to Figure 3*

**(A)** Mice received injection with AAV-CaMKII-hM3Dq-mCherry in the ACC and trained with the 6-day VI training. The first devaluation test was performed following the $4^{th}$ VI training session. After the first test, mice were re-trained with the VI schedule for one additional day, after which the second devaluation test was performed. Drug injections were conducted in a cross-over design. **(B)** Effects of ACC activation on lever pressing in the devaluation tests. N=13. **(C)** Within-subject changes in the devaluation index of **(B)**. N=13. **(D)** Mice were injected with AAV-CaMKII-hM4Di-mCherry or AAV-CaMKII-hM3Dq-mCherry in the LOFC. After the 4th VI training of the two-step trainings, the omission test was conducted. Vehicle (VEH) or CNO (0.5 mg/kg) was injected 20-min before the test. **(E)** Lever press rate during the omission test normalized by the lever press rate in the last training. VEH, N=20; CNO Di, N=16; CNO Dq, N=16. **(F)** Time to the 50% or 75% of the total presses. VEH, N=20; CNO Di, N=16; CNO Dq, N=16. **(G–I)** Relationship between the execution index and time to the 50% and 75% of the total presses in the omission test in VEH-treated group **(G)**, CNO-treated hM4Di group **(H)** and CNO-treated hM3Dq group **(I)**. (G) N=20; (H) N=16; (I) N=16. **(J–M)** Ratios of intake of sucrose and chow in the free-access sessions (Left) and numbers of nose pokes during the 5-min lever pressing sessions (Right) in mice received chemogenetic manipulation of the ACC or LOFC. Each data set corresponds to the experiments shown in Fig. 3 and Fig. S3A; chemogenetic ACC inhibition (**J**; Fig. 3A–D), ACC activation after VR+VI trainings (**K**; Fig. 3E–H), ACC activation after VI+VI trainings (**L**; Fig. S3A–C) and LOFC inhibition (**M**; Fig. 3I–L). (J) N=11; (K) N=10; (L) N=13; (N) N=13. *P < 0.05; **P < 0.01; N.S., not significant.

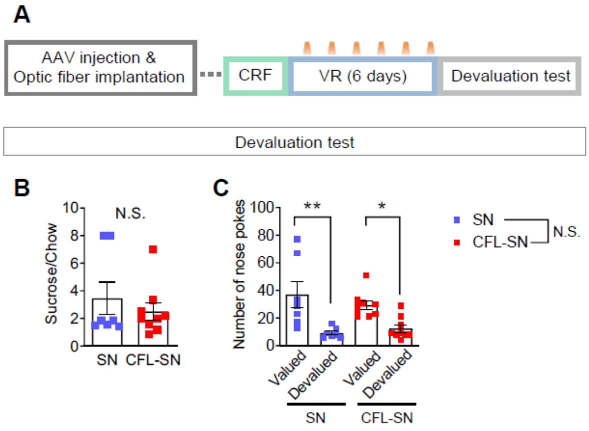


**Figure S4**

*Preference and satiety for sucrose rewards in the devaluation tests are not impaired by CALI in RSC-projecting ACC neurons, related to Figure 4*

**(A)** As shown in Fig. 4K–N, mice received CALI (illumination with 1–2 mW amber light for 1 min) in RSC-projecting ACC neurons within 5-min after each VR trainings(day 3–9), which was followed by the devaluation test. **(B, C)** Ratios of intake of sucrose and chow in the free-access sessions **(B)** and numbers of nose pokes during the 5-min lever pressing sessions **(C)**. Each data set corresponds to the experiments shown in Fig. 4M–O. SN, N=7; CFL-SN, N=9. *P < 0.05; **P < 0.01; N.S., not significant.

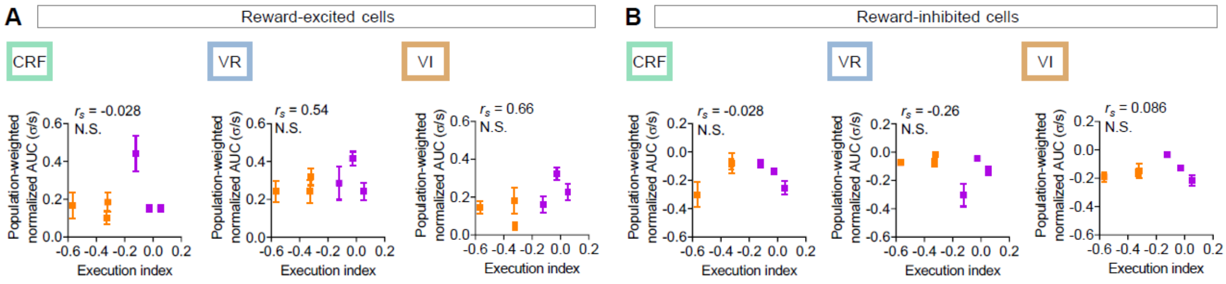


**Figure S5**

*Relationship between the execution index and reward-related responses in the RSC-projecting ACC neurons, related to Figure 5*

**(A)** Relationship between the execution index and population-weighted normalized AUC in the reward-excited cells shown in Fig.5I. CRF, N=47; VR, N=99; VR+VI, N=69. **(B)** Relationship between the execution index and population-weighted normalized AUC in the reward-inhibited cells shown in Fig.5J. CRF, N=71; VR, N=59; VR+VI, N=69. N.S., not significant.

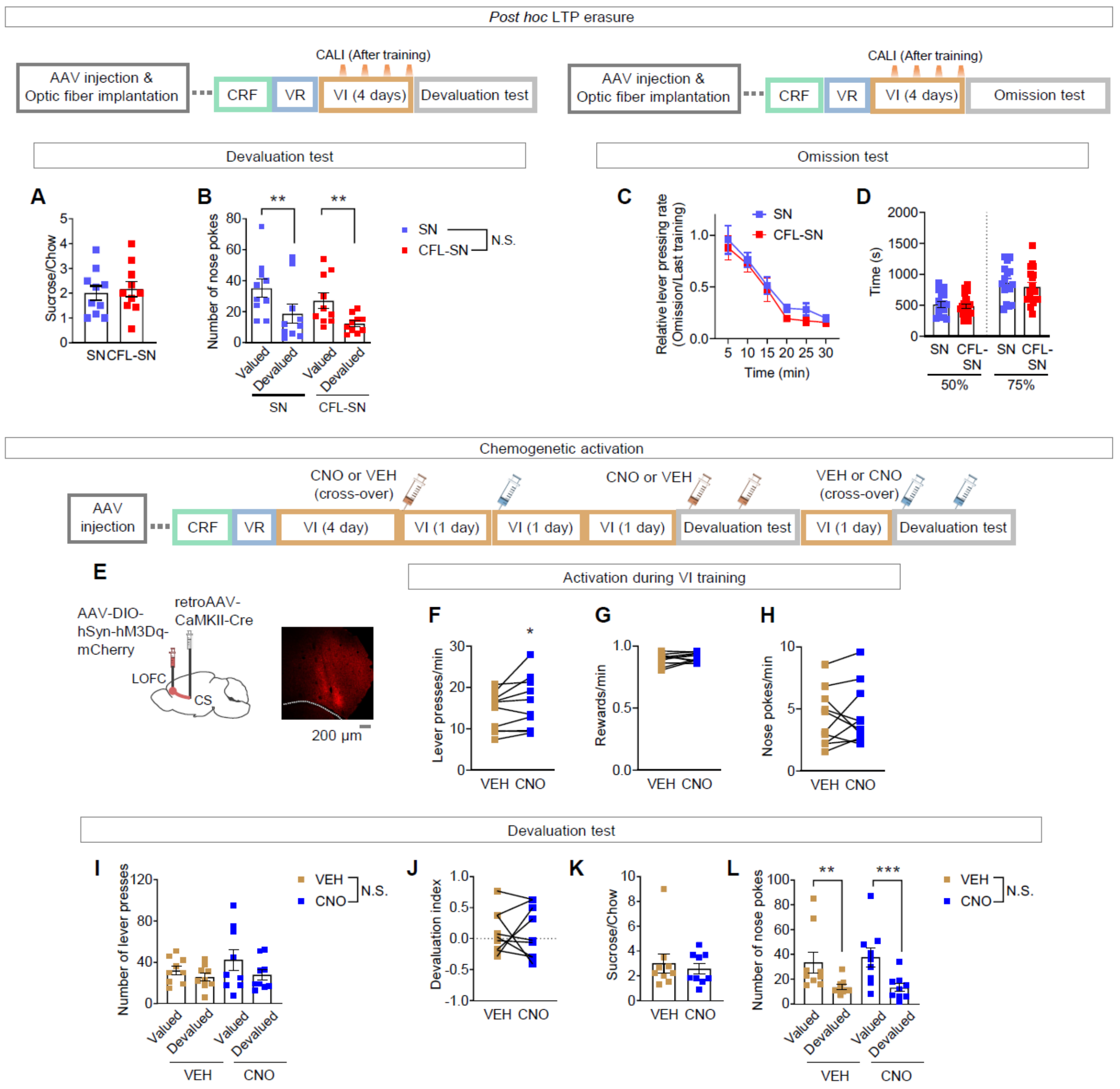


**Figure S6**

*Effects of manipulation of CS-projecting LOFC neurons on decision-making strategy and execution, related to Figure 6*

**(A–D)** Mice received CALI (illumination with 1–2 mW amber light for 1 min) in CS-projecting LOFC neurons within 5-min after each VI training (day 10–13) which is followed by devaluation **(A, B)** or omission **(C, D)** tests. Each data set corresponds to the experiments shown in Fig. 6I–M. **(A)** Ratios of intake of sucrose (g) and chow (g) in the free-access sessions. SN, N=10; CFL-SN, N=10. **(B)** Numbers of nose pokes during the 5-min lever pressing sessions. SN, N=10; CFL-SN, N=10. **(C)** Lever press rate during the omission test normalized by the lever press rate in the last training. SN, N=16; CFL-SN, N=19. **(D)** Time to the 50% or 75% of the total presses. SN, N=16; CFL-SN, N=19. **(E)** Mice were injected with AAV-DIO-hSyn-hM4Di-mCherry in the LOFC and retroAAV-CaMKII-hM3Dq-mCherry in the CS. **(F–H)** After the 4th VI training of the two-step trainings, mice were trained two additional VI sessions with injection of CNO or its vehicle in a cross-over design. **(F–H)** Effects of CNO on rates of lever pressing **(F)**, reward presentations **(G)** and nose pokes **(H)** during the VI session. N=10. **(I–L)** After experiments shown in **(F–H)**, mice were re-trained with a one-day VI session, which was followed by devaluation tests with injection of CNO or its vehicle in a cross-over design. **(I,J)** Number of lever presses **(I)** and devaluation index **(J)** in each devaluation test. **(K,L)** Ratios of intake of sucrose and chow in the free-access sessions **(K)** and numbers of nose pokes during the 5-min lever pressing sessions **(L)**. N=9. *P < 0.05; **P < 0.01; ***P < 0.001; N.S., not significant.

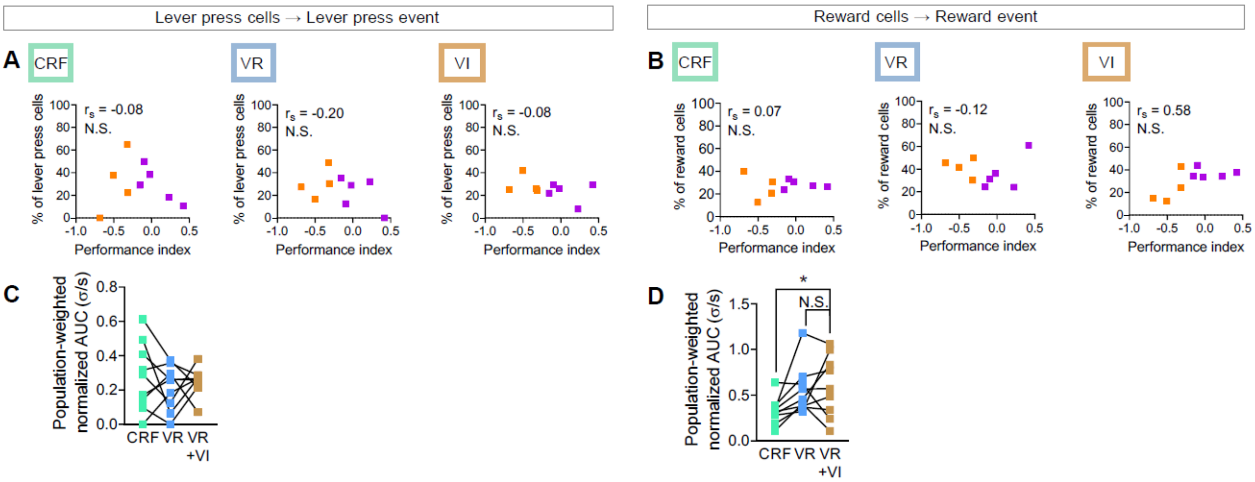


**Figure S7**

*Relationship between the execution index and fractions of the event-related cells in the CS-projecting LOFC neurons, related to Figure 7*

**(A)** Relationship between the execution index and fractions of the lever press cells for each mouse shown in Fig.7G & H. **(B)** Relationship between the execution index and fractions of the reward cells for each mouse shown in Fig.7I & J. **(C)** Within-subject comparisons of population-weighted AUCs of the lever press cells to lever press events during each training stage. **(D)** Within-subject comparisons of population-weighted AUCs of the reward cells to reward events during each training stage. N=9 mice *P < 0.05; N.S., not significant.
